# Evidence integration and decision-confidence are modulated by stimulus consistency

**DOI:** 10.1101/2020.10.12.335943

**Authors:** Moshe Glickman, Rani Moran, Marius Usher

## Abstract

Evidence-integration is a normative algorithm for choosing between alternatives with noisy evidence, which has been successful in accounting for a vast amount of behavioral and neural data. However, this mechanism has been challenged as tracking integration boundaries sub-serving choice has proven elusive. Here we first show that the decision boundary can be monitored using a novel, model-free behavioral method, termed *Decision-Classification Boundary*. This method allowed us to both provide direct support for evidence-integration contributions and to identify a novel integration-bias, whereby incoming evidence is modulated based on its consistency with evidence from preceding time-frames. This consistency bias was supported in three cross-domain experiments, involving decisions with perceptual and numerical evidence, which showed that choice-accuracy and decision confidence are modulated by stimulus consistency. Strikingly, despite its seeming sub-optimality, this bias fosters performance by enhancing robustness to integration noise. We argue this bias constitutes a new form of micro-level, within-trial, confirmation bias and discuss implications to broad aspects of decision making.

## Introduction

Evidence integration to boundary is a normative mechanism for evidence-based decisions, which provides the fastest mean response-time (RT) for a target accuracy rate (Bogacz, Brown, Moehlis, Holmes, & Cohen, 2006; Gold & Shadlen, 2002; Moran, 2015; Wald, 1947) and accounts for an impressive amount of behavioral and neural choice data (see Forstmann, Ratcliff, & Wagenmakers, 2016 for a review). For example, integration-to-boundary models (Bocacz et al., 2006; Brown & Heathcote, 2008; Gold & Shadlen, 2001; Ratcliff & McKoon, 2008; Teodorescu & Usher, 2013; Usher & McClelland, 2001; Vickers, 1970) provide a parsimonious account for the shape of choice-RT distributions of correct and incorrect responses as a function of stimulus difficulty, as well as for the well-known speed accuracy trade-off (Wickelgren, 1977). Moreover, integration-to-boundary models are supported by the monitoring of neural activation in brain decision areas during choice tasks (Gold & Shadlen, 2007; Mulder, van Maanen, & Forstmann, 2014; but see Latimer, Yates, Meister, Huk, & Pillow, 2015).

Despite this strong support, however, in behavioral studies the decision-boundary remains a theoretical concept, which is usually inferred from the data via model fitting and depends on specific model assumptions. Moreover, this framework can be challenged by alternative non-integration mechanisms, such as heuristics based on the detection of a single high-value sample, which can account for many of these choice-patterns (Ditterich, 2006; Watson, 1979; see also discussion in Stine, Zylberberg, Ditterich, & Shadlen, 2020). It is thus imperative to validate the *integration* assumption and to directly monitor the decision boundary. While one can support integration to boundary via quantitative model comparison (Stine et al., 2020), such comparisons are limited by specific and somewhat arbitrary parametric assumptions (e.g., a particular temporal collapsing regimen of a choice threshold). Furthermore, recent findings indicate a variety of biases in the sampling and weighting of evidence (Bronfman et al., 2015; Cheadle et al., 2014; Dotan, Meyniel, & Dehaene, 2018; Glickman, Tesetsos & Usher, 2018; Glickman et al., 2019; Gluth, Kern, Kortmann, & Vitali, 2020; Rollwage et al., 2020; Talluri et al., 2018; Urai, de Gee, Tsetsos & Donner, 2019). Potential biases of this kind should be taken into account when assessing whether evidence is accumulated. Here, we addressed these issues using a novel behavioral and model-agnostic method, which we term *Decision-Classification Boundary* (DCB).

When evidence integration is perfect (i.e., free of distortions and biases), the DCB recovers the decision boundary. More broadly, however, DCBs provide a novel behavioral signature — a benchmark for evaluating biases in evidence-integration. Applying this method to data from three experiments across choice domains (numerical cognition and perception), we find strong support for evidence-integration over heuristic non-integration models. Furthermore, we demonstrate an important new factor modulating evidence accumulation – *stimulus-consistency*, corresponding to an increased relative weighting of pieces of evidence preceded by information supporting the same choice-alternative, resulting in a type of *momentary confirmation-bias (*Bronfman et al., 2015; Rollwage et al., 2020; Talluri et al., 2018), which operates during (rather than after) a decision. Importantly, this mechanism achieves an important computational advantage: it makes the decision robust to late (non-encoding) noise (Spitzer et al., 2017; Tsetsos et al., 2016).

We start with a description of our experimental design (showing it is possible to extract behavioral signatures of integration to boundary), followed by a computational section that presents the model-free DCB method. We then apply this method to the data from two experiments, focusing on the dependency of the DCB on stimulus consistency and resort to computational modeling to specify the stimulus-consistency mechanism. Finally, we present the results of a third experiment designed to test this mechanism.

## Results

### Experimental design and behavioral signature of integration to boundary

In two experiments (Exp. 1, reported in Glickman & Usher, 2019 and Exp. 2, novel data), participants were presented with sequences of pairs of stimuli (two digit numbers or bars, respectively; see Fig. 1A-B), which were sampled from two overlapping Normal distributions (see Experimental Methods). The sequences were presented at a rate of 2 pairs/sec (numbers) and 5 pairs/sec (bars), and were terminated by the participant’s response. The task was to select the sequence that corresponded to the higher generating distribution (red Gaussian in Fig 1A-B). Between trials, we manipulated stimulus difficulty by varying the separation between the Normal distributions (see Experimental Method for further details). Twenty-seven participants performed 500 trials in Exp.1 and thirty participants performed 480 trials in Exp.2.

**Figure 1.**
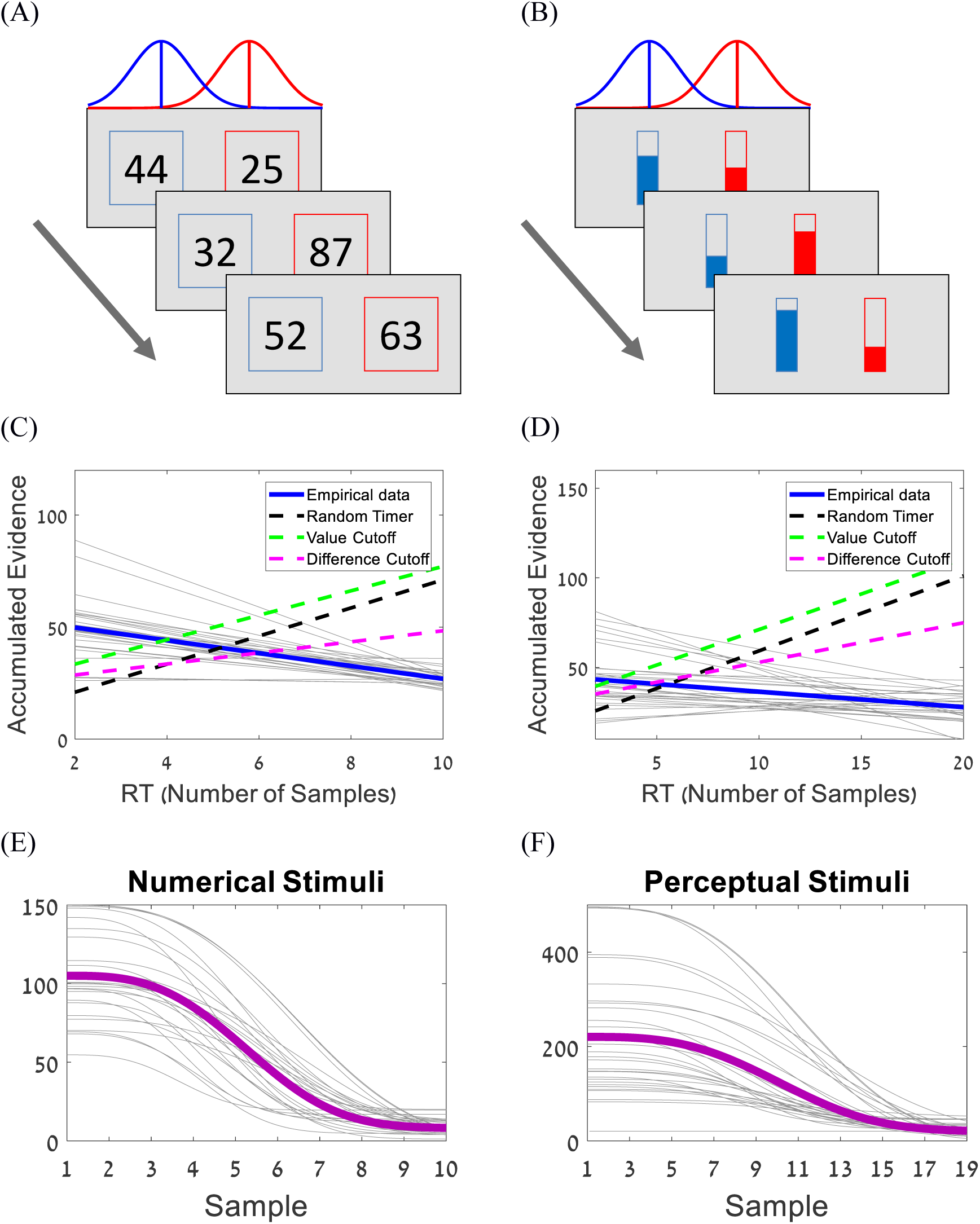
(A-B) Experimental paradigms; participants are presented with pairs of numerical values (Exp. 1) or bars (Exp. 2) sampled from two overlapping Normal distributions, and are asked to choose which sequence was drawn from a distribution with a higher mean. The presentation is terminated by the decision of the participant. (C) The accumulated evidence of all participants (blue line) as a function of decision-time; grey lines correspond to accumulated evidence of individual participants. The black, green and magenta dashed lines correspond to the random time model, and to the value and difference heuristics, respectively. (D) Same as (C) for Exp. 2. (E-F) The collapsing-boundaries obtained in Exp. 1 (E) and 2 (F); Solid purple lines correspond to the boundaries generated using the group mean parameters, grey lines correspond to the boundaries of the individual participants.

In previous work, we showed that the choices in Exp. 1 (numerical evidence) support integration of evidence to a collapsing boundary and excluded a set of non-integration models (Glickman & Usher, 2019). Here, we extend this analysis to the data in the perceptual domain (Exp. 2). The blue lines (group data) and thin grey lines (individual subjects) in Fig 1 C-D are obtained by integrating the trial-by-trial stimulus evidence until the decision moment (i.e., the cumulative sum of differences between the sequence), and averaging across trials for each RT (see Glickman & Usher, 2019 and Suppl. A for further details). These lines show a mildly decreasing pattern (Fig 1C: *b* = −2.86, *t* = −7.08, *p* <. 001, and Fig 1D: *b* = −0.86, *t* = −3.18, *p* =. 001), which is the behavioral signature of integration to a collapsing boundary (see Suppl. A). These results are consistent with model fitting using a Weibull parametrization of the boundary (Hawkins et al., 2015), which also indicates a collapsing boundary (Fig. 1E-F). Critically, the descending slopes of integrated evidence rule out non-integration strategies, such as: i) random-timer – response time is determined by a process that is exogenous to the integration of evidence (black dashed line), ii) value-cutoff – observers choose the sequence in which a number exceeding some predetermined threshold first appears (green dashed line), and iii) difference-cutoff – observers choose based on the first frame in which the difference between the numbers exceeds a predetermined threshold (magenta dashed line). All of these non-integration models predict that the integrated evidence increases (rather than decreases) with the number of samples (see Computational Methods for further details about these strategies). This is because, if the stopping rule is independent of the evidence integrated so far, longer decision trials necessarily accumulate more evidence. While these results provide support for integration to boundary, the actual shape of the boundary trajectory (Fig. 1E-F) is only extracted via model fitting, as the evidence-integration lines (Fig 1 C-D, blue and thin grey lines) are systematically biased by the accumulation of noise during the trial. In the following, we present a novel model-free method to estimate the decision boundary via a classification boundary curve, which is more robust to accumulation noise and reconstructs the actual shape of the decision boundary.

### Model-Free extraction of decision boundaries: the boundary classification curve

We simulated synthetic data based on the experimental task we used in our experiments (see Fig. 1A-B and Computational Methods), in which sequences of values are sampled from two overlapping Normal distributions. It is assumed that the subjects integrate a noisy version of the evidence at each frame, according to the following difference equation:

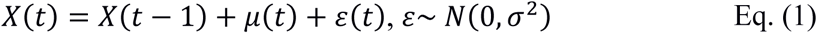

where *X(t)* is the accumulated differences between the sequences at time *t*, *μ*(*t*) is the difference between the samples at time *t*, and ε(*t*) is a temporally-independent random internal Gaussian noise generated by the encoding system, which is independent from the evidence-sampling noise (Fig 1A-B). We also assumed that a response is triggered when the integrated noisy evidence reaches one of two symmetric decision boundaries. Two types of boundaries were used in these simulations: fixed and collapsing (Hawkins et al., 2015). For the collapsing boundary, we follow Hawkins et al. (2015) to parameterize the boundary using Weibull functions (see Computational Methods). In each simulation, we ran 10,000 trials, in which (after sampling values from the two distributions and integrating subject to noise), we record the choice, as well as the input sequences (without the internal noise) sampled until the decision was made.

Based on this data, we reconstruct the boundary at each time frame using a method based on Linear Discriminant Analysis (LDA; Fisher, 1936; McLachlan, 2004), which generates boundary classification curves. These curves are obtained by applying LDA to the integrated-evidence excluding the random internal noise, defined as:

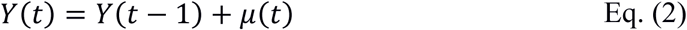

The LDA was applied to the integrated-evidence (*Y(t)*) of the observer at each time frame, across all trials in the experiment, so as to classify the action at each time frame to one of three categories: choose alternative A (Fig. 2A-B, blue distributions), choose alternative B (Fig. 2A-B, green distributions), or continue sampling (Fig. 2A-B, orange distributions). The classification boundary curve (Fig. 2A-B, red line) best separates the different classes (see Computational Methods). Note that in the absence of internal noise, the true decision boundary would separate perfectly these categories. However, in the presence of internal noise, which may increase the values of integrated evidence that lie below the boundary (blue areas), or vice versa (orange areas above the boundary), there is some unavoidable overlap.

**Figure 2.**
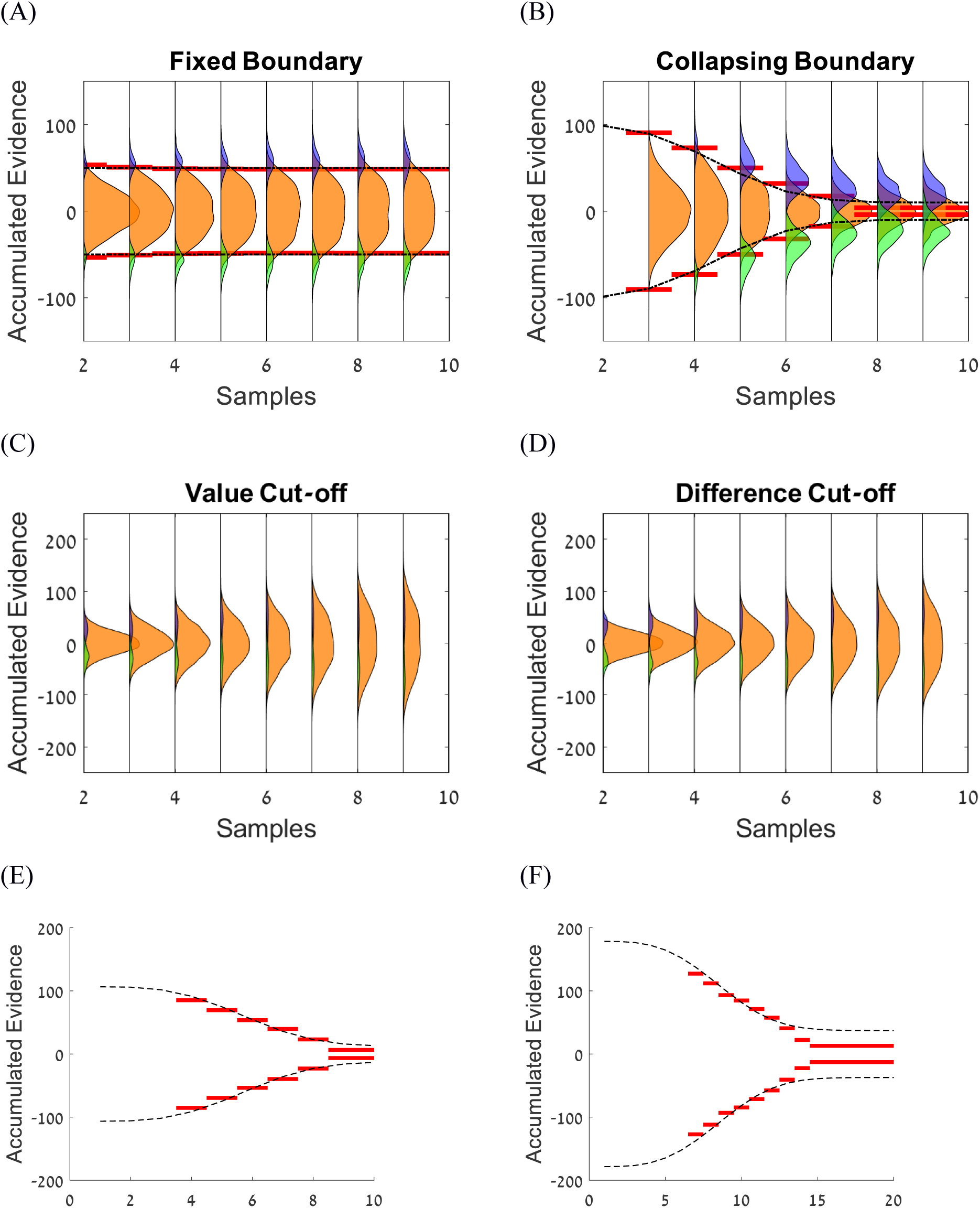
(A-B) Illustration of the boundary extraction for a decision simulated using a Diffusion model with fixed (A) or collapsing (B) boundary. The black dashed line corresponds to the original boundary with which the model was simulated and the red lines correspond to the modelfree best-fitted DCB. The orange distributions correspond to data points within each trial in which the simulated participant continued sampling, while the blue and green distributions correspond to frames in which the decision was terminated. The total area under the orange, blue, and green distributions was normalized for each frame. (E-F) Experimental data of a representative subject that participated in the numerical (E) and perceptual (F) experiments. Black dashed line corresponds to the model-based best-fitted boundaries, and the red lines correspond to the model-free best-fitted boundaries of the experiments, respectively.

The results are illustrated in Fig. 2A-B, which shows that the DCBs (red solid lines) recover quite accurately the generating boundaries (black dashed lines). In particular, the extracted boundary is temporally constant or decreases as a function of time, when the generating boundary is flat or collapsing, respectively. Notably, the quantitative agreement between the generating and recovered boundaries is high. Note that the model-free boundary extraction method makes no *a priori* parametric assumptions about the shape of the model boundary. As shown in Fig. 2C-D, for synthetic data generated using the non-integration-to-boundary value (or difference) cutoff heuristics, the three distributions of evidence – trials in which the model continues sampling (orange) and trials in which response has been made (blue and green) – totally overlap. Consequently, the decision classification curve algorithm fails to classify the three classes of trials based on integrated evidence. Thus, the presence of stable boundary classification curves (Fig. 1A-B) provides strong support against (non-integration) cutoff models. In both our data sets, we find stable classification curves (see Fig. 3E-F, red lines, for representative example subjects) that imply a collapsing boundary, which are consistent with model fitting (Fig. 3E-F, black dotted line), but make no parametric assumption on the form of this boundary (see Suppl. C for DCB of all participants).

**Figure 3.**
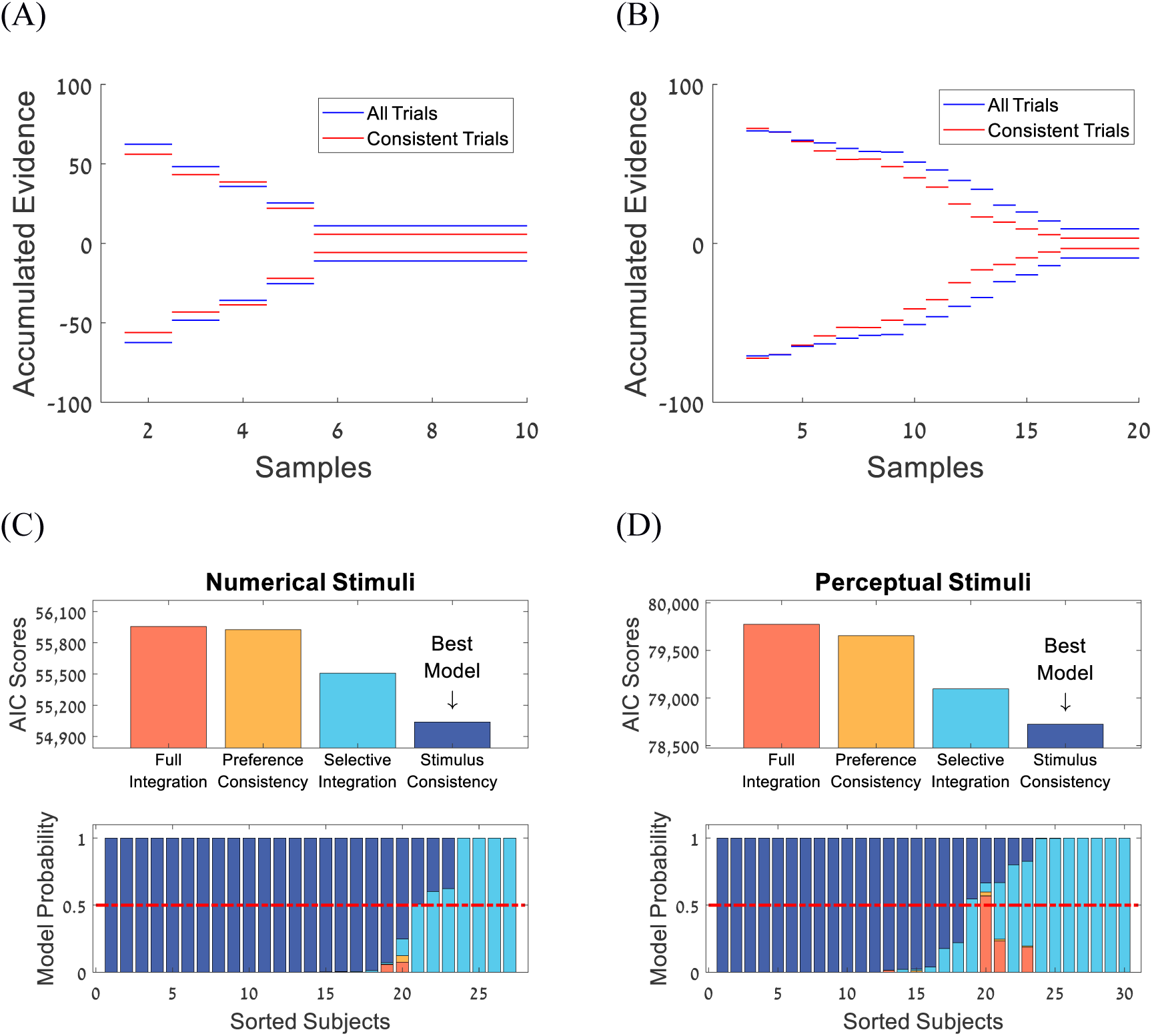
Results of Exp. 1 & 2. (A-B) Decision boundaries for all trials (blue) and high consistency trials (red) of representative participants in (A) Exp. 1 - numerical stimuli and (B) Exp. 2 - perceptual stimuli. (C-D) The stimulus-consistency model outperformed the full integration, preference consistency and selective integration models in Exp. 1 - numerical stimuli (C), as well as in Exp. 2 - perceptual stimuli (D).

### Evidence integration is modulated by stimulus consistency

A more detailed examination of the choice data in both experiments shows that *stimulus consistency* – operationalized as the absolute value of the difference between the number of frames with evidence favoring each alternative divided by the total number of frames – has a critical impact on choices and RT, above and beyond the effect of total evidence. To illustrate, consider two trials with the same total evidence: Trial 1: (2, 3, 1, 4, total evidence = 10) and Trial 2: (6, −1, 8, −3, total evidence = 10). While both trials have a total evidence of 10 in favor of one of the alternatives, the evidence stream is more consistent in Trial 1 (consistency measure 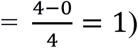) compared with Trial 2, where half of the evidence favoring one alternative whereas the other half favors the other (consistency measure 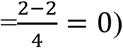). Previous research has shown that stimulus consistency modulates decision confidence (Dotan et al., 2018). Here, we examine its impact also on accuracy, RT, as well as on the shape of decision boundaries. To this end, we conducted several mixed-model regression analyses (logistic for accuracy and linear for RT and confidence), in which we predicted trial by trial choice accuracy, RT, and confidence, using accumulated evidence and stimulus consistency as fixed factors and participants as random intercepts. The results, which are shown in Table 1, indicate that stimulus consistency improves accuracy and confidence and reduces RT, independent of the accumulated evidence.

**Table 1.** Beta coefficients for predicting Accuracy, RT, and confidence in Exp. 1&2.

|  | **Exp. 1**<br>**(Numerical Stimuli)** |  | **Exp. 2**<br>**(Perceptual Stimuli)** |  |  |
| --- | --- | --- | --- | --- | --- |
|  | **Accuracy** | **RT** | **Accuracy** | **RT** | **Confidence** |
| **Evidence** | $\beta = 1.71^{***}$<br>$t = 27.90$ | $\beta = 0.26^{***}$<br>$t = -28.21$ | $\beta = 0.55^{***}$<br>$t = 16.91$ | $\beta = -0.16^{***}$<br>$t = -20.42$ | $\beta = 0.13^{***}$<br>$t = 12.14$ |
| **Stimulus Consistency** | $\beta = 0.26^{***}$<br>$t = 5.69$ | $\beta = -0.13^{***}$<br>$t = -14.41$ | $\beta = 0.27^{***}$<br>$t = -9.40$ | $\beta = -0.13^{***}$<br>$t = -16.78$ | $\beta = 0.14^{***}$<br>$t = 13.96$ |

\*\*\* $p < .001$

Note that the "Difference in Evidence Directions" (DED) is only one measure of stimulus consistency. More complex forms that include temporal factors can also be construed. For example, the consistency bias may correspond to the size of the larger temporal cluster (LTC) with evidence in the same direction (i.e., the largest cluster of evidence; see Suppl. B for analysis showing that such an LTC-measure also predicts differences in accuracy, RT, and confidence independent from total evidence).

### Decision classification curves modulation by stimulus consistency and model comparison

Motivated by the results above, we examined if stimulus-consistency modulates the decision classification curve. Towards this aim, we extracted the DCB of high consistency trials (DED measure higher than 0.5) using the model-free method, and compared it with the DCB of all trials, which correspond to the most accurate estimation of the boundary (in the absence of a consistency-modulation). As illustrated below (Fig. 3 A-B; for two representative subjects; see Suppl. C for all subjects), in both experiments, the DCB is lower for the high consistency trials (Fig. 3 A-B; red lines) than for all trials (Fig. 3 A-B; blue lines; Exp. 1: *t*(26) = 2.67, *p* =. 01, Exp. 2: *t*(29) = 3.91, *p* <. 001; see Suppl. D for further details). Critically, we tested that applying the same boundary reconstruction procedure to synthetic data simulated from a full integration model, does not result in DCB differences based on the consistency of the evidence (see Suppl. D).

These findings could suggest either that under high stimulus consistency the participants require less evidence to commit to a decision (i.e., choice boundaries are truly lower) or alternatively, that evidence integration itself may be modulated towards the same decision boundary (for high consistency trials); an evidence sample might be boosted when it comes after another sample supporting the same alternative, but evidence is integrated to the same boundary. As our interest is on the functional mechanism, we do not aim to distinguish these two variants here. Rather, we see the DCB as an empirical measure (which is data and not model-driven), against which models can be evaluated.

To specify the evidence-integration mechanism, we carried out a quantitative model comparison on the likelihood to account for the data (response and number of samples taken) for each trial of each participant in the two experiments (we used AIC as a measure of fit to include a model-complexity cost). We started with the perfect integration model (which assumes no systematic distortion of the evidence a subject integrates in each frame, i.e., integration is only corrupted by additive noise). Next, we examined a selective-integration (SI) mechanism that amplifies or diminishes, respectively, the stronger or weaker evidence within each frame across the two evidence streams (Tsetsos et al., 2016; Usher, Tsetsos, Glickman, & Chater, 2020; see Computational Methods). Critically, we also examined several variants of a “stimulus-consistency” model. In the simplest variant, the evidence is modulated solely based on whether evidence from the preceding frame is consistent (i.e., it points in the same direction) with the current frame. In a slightly more complex variant, the modulation magnitude increases linearly with the number of consistent frames and resets to baseline (no modulation) with every swap (we report here only the results of the more complex version, which provided better fit for the data). Finally, we also examined a *preference-consistency* model, wherein incoming evidence is modulated based on consistency with the total integrated evidence (thus reflecting momentary preference) up to that time point (see Computational Methods for details).

In each model, we allowed for collapsing boundaries, which, as illustrated in Fig 2C-D, capture well the shape of the decision boundary (and provide much better fits to the data compared to fixed boundaries; Glickman & Usher, 2019). Fig. 3C-D shows group and individual participant fit measures for four models: full integration, selective-integration, preference-consistency, and stimulus-consistency in Exp. 1 & 2. As illustrated, the most successful of the models was the stimulus-consistency version in which evidence increased at each consecutive frame in the same direction, followed by the SI-model.

### Experiment-3: testing the stimulus-consistency effects

Because in our first two experiments, participants’ choices terminated the information stream, trial-consistency depended, at least partially, on participants’ responses. To address this limitation, and to directly test the impact of stimulus consistency on evidence-based choice and on decision confidence, a third experiment was designed. In this experiment, the number of samples was fixed, and the stimulus consistency was manipulated independently from the total evidence. Sequences of eight number pairs were presented at a rate of 2/sec (as in Exp. 1) and participants were instructed to choose (at the end of the presentation) the higher average sequence. For each set of eight value-pairs, we generated two yoked trials, which consisted of the very same evidence content (across the eight time frames) for each sequence and differed only in temporal order of the values (i.e., the yoked trials varied in how the values on each side were shuffled). In the “consistent” trial condition (Fig. 4A, bottom panel), one evidence stream provided stronger evidence in 7 out of the 8 frames, whereas in the “inconsistent” trial condition (Fig. 4A, upper panel), each stream provided stronger evidence in 4 frames.

**Figure 4.**
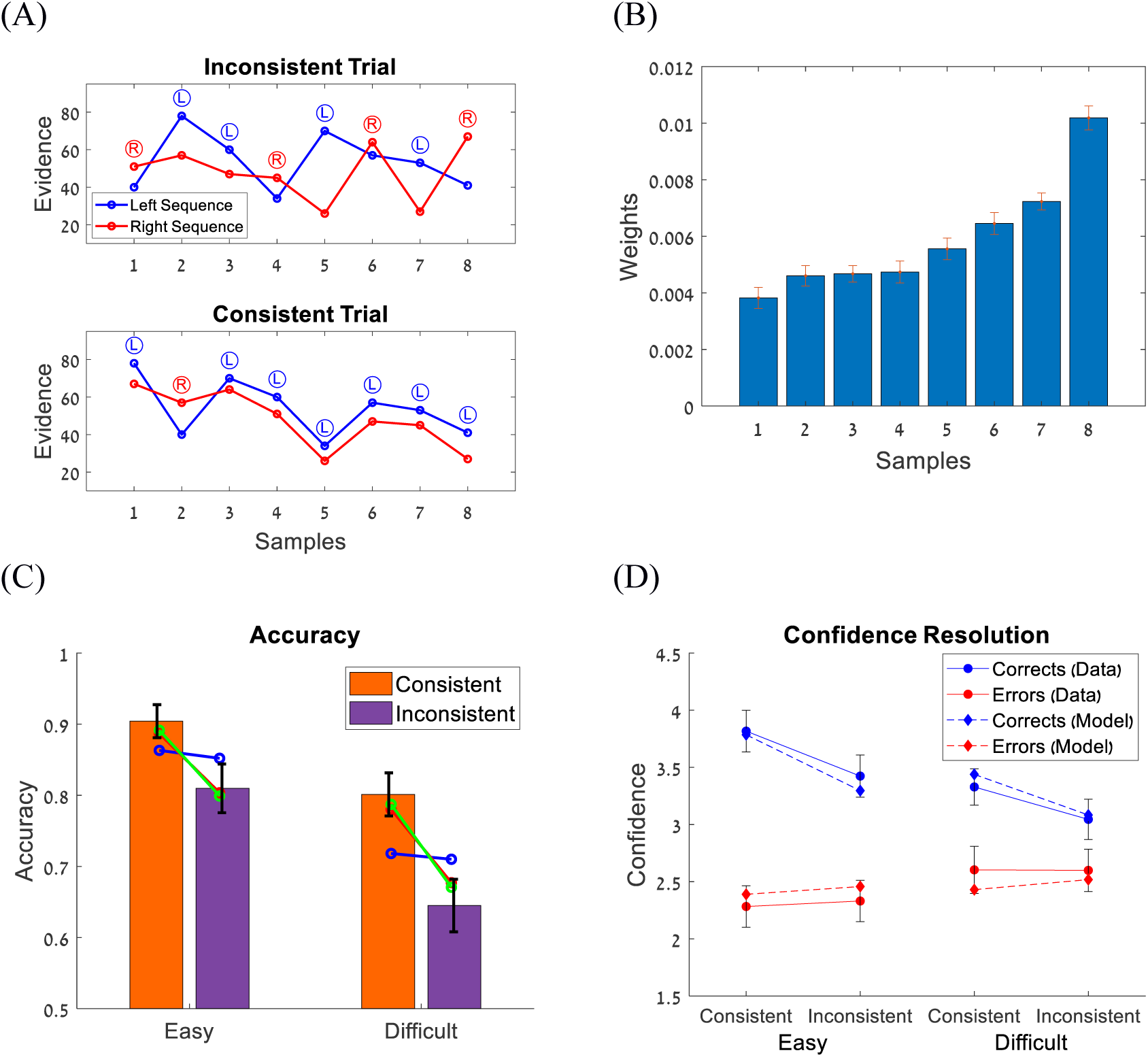
The results of Exp. 3. (A) Illustration of consistent and inconsistent trials. Note that both trials have the same amount of evidence. L and R symbols correspond to momentary advantages of the left or right sequences, respectively. (B) The impact of the temporal position (evidence) on choice was assessed using logistic regression (see Computational Methods), and revealed a recency effect, which necessitated the use of a leak term in the model. (C) Choice accuracy as a function of difficulty (separation between the sampling distributions: easy vs. difficult) and consistency (difference in the directions of the evidence: consistent vs. inconsistent). The blue, red and green lines corresponds to the predictions of the leaky-integration, selective integration and stimulus consistency models, respectively. (D) Confidence as a function of difficulty and consistency for correct (blue lines) and incorrect (red lines) responses. Data are shown with solid lines and circle symbols and model predictions are shown with dashed lines and diamond symbols. Error bars are within-subjects SE.

As shown in Fig. 4C, participants were more accurate in consistent than inconsistent trials, both in the easy, *t*(21) = 6.15, *p* <. 001, and difficult conditions, *t*(21) = 6.90, *p* <. 001. A similar pattern was also obtained for confidence, *t*(21) = 4.76, *p* <. 001, for the easy condition and *t*(21) = 4.19, *p* <. 001, for the difficult condition (Fig. 4D). Interestingly, the modulation of the confidence responses was different for correct and incorrect responses. Whereas for correct responses, confidence increases with stimulus consistency, *t*(21) = 4.05, *p* <. 001 and *t*(21) = 3.59, *p* =. 002, for the easy and difficult conditions, respectively, this pattern was not obtained for incorrect trials, *t*(21) = −0.64, *p* =. 52 and *t*(21) =. 06, *p* =. 95. These findings indicate that metacognitive accuracy (confidence-resolution = confidencecorrects – confidenceerrors) increases with stimulus consistency, both for easy trials: *t*(21) = 3.68, *p* =. 002 and difficult trials: *t*(21) = 2.59, *p* =. 02. As we show in Suppl. F, this effect can also be accounted for by the stimulus-consistency model by applying a signal-detection confidence approach.

We conducted quantitative model comparison for three types of models using the data from Exp. 3. Since the data also shows a recency pattern (see Fig. 4B and Computational Methods), we used leaky-integration (Ossmy, Moran, Donner & Usher, 2013; Roe et al., 2001; Usher & Mclelland, 2001) instead of full integration as our default model (with leak as a model parameter, accounting for decaying temporal weights). We compared a leaky-integration model (in which there is no distortion of the integrated evidence, other than the decaying temporal weights), an SI-model (which, additional to integration leak, gives higher weight to high values compared to low values, within each frame) and a stimulus-consistency model (from Exp. 1-2; Fig 3C-D). The results show that the stimulus-consistency model outperforms the full and selective integration models, and provides the best account for the data (Fig. 5).

**Figure 5.**
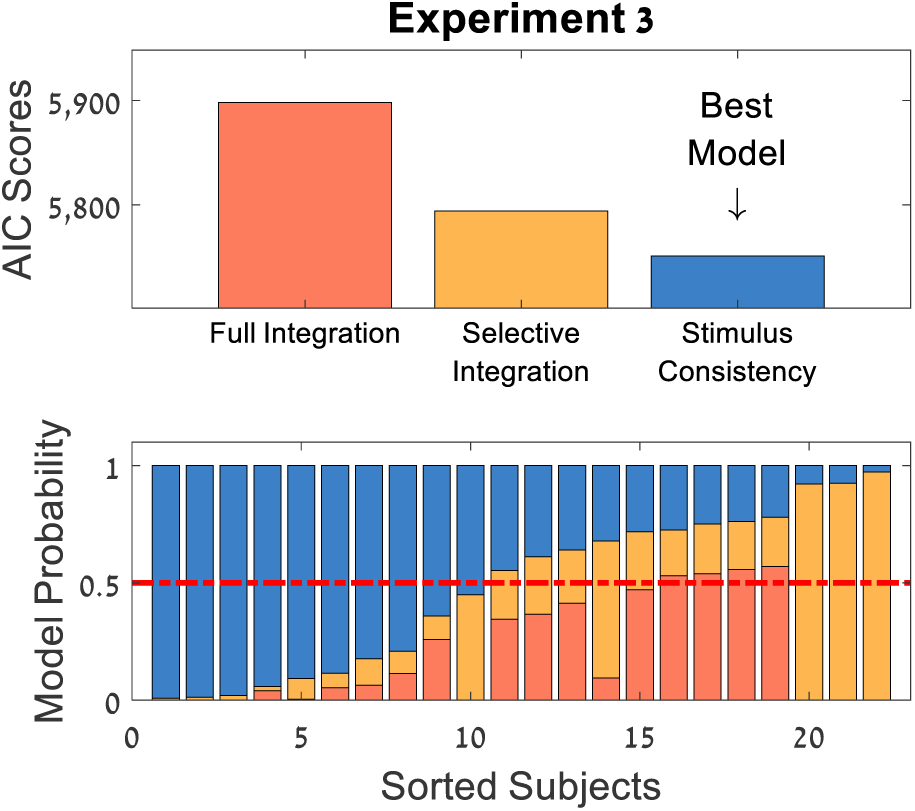
Model Comparison for Exp. 3. The stimulus consistency model outperformed the leaky and selective integration models.

Interestingly, whereas both of the bias models are able to account for the modulation of accuracy by stimulus consistency in Fig 4C, the stimulus consistency bias model accounts for subtler temporal clustering effects in the data. For example, unlike the SI model, the stimulus-consistency model predicts that accuracy is modulated by the size of the largest cluster of evidence consistent with the correct choice (LTC; see Suppl. B for the impact of this measure in Exp. 1 & 2 and Suppl. E for data showing an association between the LTC-enhancement and the advantage of stimulus-consistency over the SI model in Exp. 3).

### Stimulus-consistency and normativity

#### Normative considerations

Why do participants increase the relative weighting of pieces of evidence that are consistent with previous ones? At face value, this distortion introduces a gap between the accumulated and ‘real’ evidence and should reduce task performance. Indeed, this is the case in the absence of (or for low) integration noise. As illustrated in Fig, 6A-B, however, in the presence of high integration noise, the consistency modulation makes the mechanism more robust to the corrupting impact of this noise^1^ (see cross-over between red and light blue lines in Fig 6A, so that for each level of noise, there is a consistency-modulation that optimizes performance; black dots in Fig 6B and dark blue line in Fig 6A). A similar robustness effect due to selective-integration was reported by Tsestos et al. (2016) for the SI-model (see also Spitzer et al., 2017 and Usher, Tsestsos, Glickman & Chater, 2019). In both cases, the integration mechanism distorts the actual evidence by shifting the distribution of accumulated evidence towards the correct side (see center of the blue/red Gaussians in Fig 4C-D, while also making this distribution broader). While for low noise this is detrimental to performance, for high noise it is beneficial, as the shift helps to make the effect of additional noise less pronounced.

**Figure 6.**
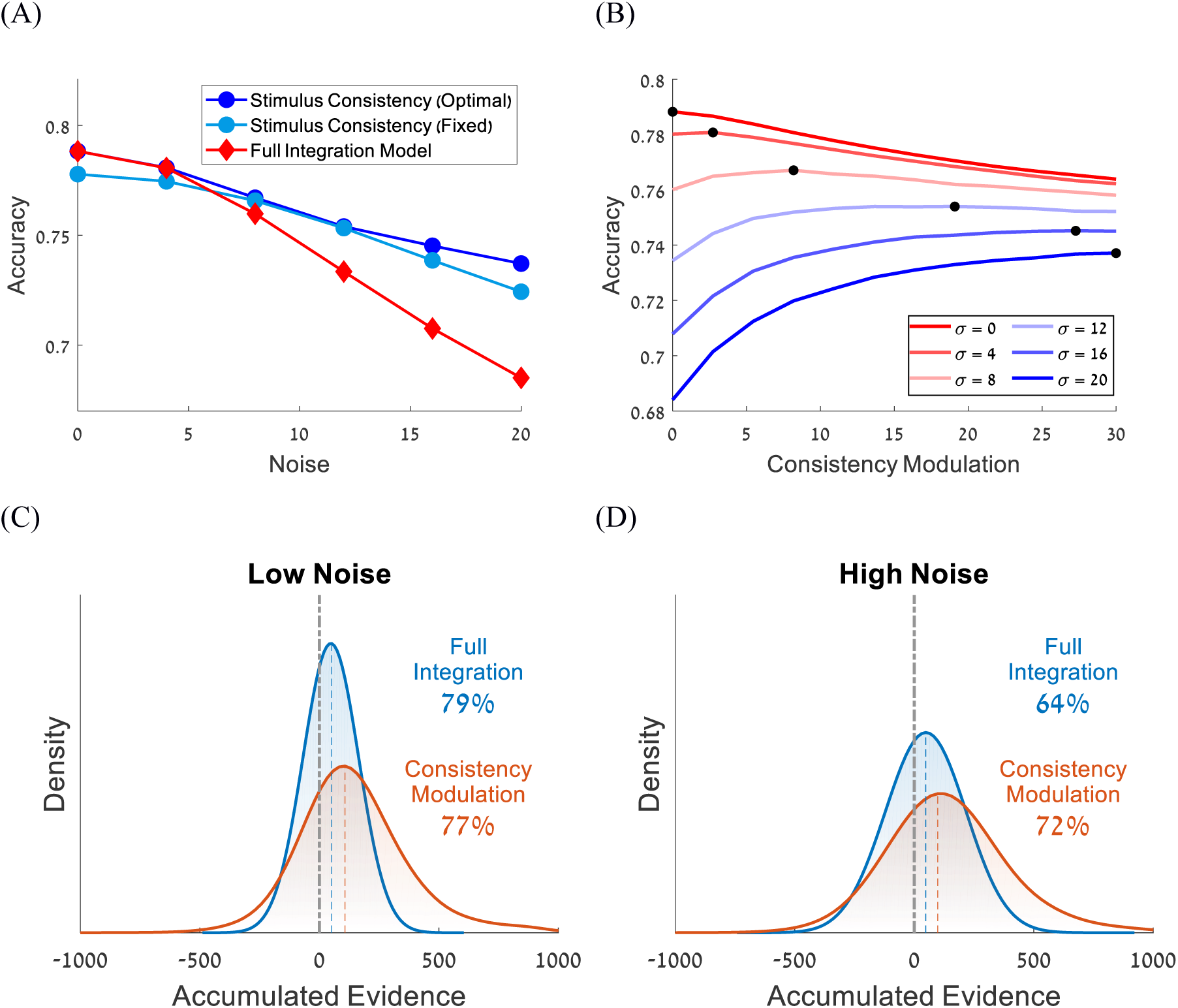
Consistency based modulation and normativity (A) accuracy (i.e., robustness to noise) as a function of noise, separately for the full integration (red line) and stimulus consistency models (blue lines). The accuracy of the stimulus-consistency model is presented for the model simulated using a fixed consistency parameter value of 10 (light blue) and for the model simulated using the optimal consistency parameter for each noise level (dark blue; see also panel B). (B) Accuracy of the stimulus-consistency model as a function of the consistency-modulation strength, for different levels of noise (σ curves). Black circles indicate consistency-values that maximize accuracy for a given level of accumulation noise. One can see that as the level of the noise increases (blue to red lines), so does the level of the consistency modulation needed to achieve optimal performance (black circles). (C-D) The distributions of the total accumulated evidence at the moment of response for the full integration model (blue) and stimulus-consistency model (red). The accuracy of the full integration model is higher than that of the stimulus-consistency model for low noise simulations (C), however the stimulus-consistency model is more accurate for high noise simulations (D).

## Discussion

In the present study we examined the mechanism that human observers deploy when making decisions on rapid streams of stochastic evidence (perceptual and numerical). Using a novel behavioral model-agnostic method — the DCB curve – we showed that decision making is well characterized by integration to boundary rather than by non-integration heuristics, and that the boundary collapses with the passage of time (Fig. 2E-F; see also Balsdon, Wyart & Mamassian, 2019). Furthermore, we found that DCB curves constitute an informative behavioral benchmark for evaluating biases in evidence-integration. Strikingly, the full integration model failed to account for the lowered DCB for high consistency trials, indicating that integration of evidence was subject to systematic biases.

Motivated by previous studies, we focused on two types of evidence integration biases. The first is an attention bias, which affects the relative weight of evidence given to temporally simultaneous sources of evidence (Krajbich et al., 2010). In the selective-integration model, for example, on each frame the higher of the two values receives a higher weight than the lower one (Glickman et al., 2018; Luyckx et al., 2020; Tsetsos et al., 2012). The second bias involves the sequential impact of a frame on subsequent frames, whereby evidence that is consistent with previous frames receives higher weight than inconsistent evidence. Formal model comparisons, as well as analysis based on qualitative behavioral patterns, supported the superiority of consistency biases in accounting for our data. Notably, consistency affected not only decision accuracy, but also decision confidence (see also Dotan et al., 2018), such that consistent evidence facilitated high decision confidence even after controlling for the total amount of evidence. Notably, consistency did not merely exert a biasing influence on confidence but improved participant’s meta-cognitive performance as measured by the resolution of confidence (i.e., the correlation between confidence and choice correctness). Indeed, confidence as a function of consistency increased for correct choices but remained constant for incorrect choices. An open question for future studies is whether the effects that consistency exerts on choice accuracy and meta-cognition are dissociable.

Previous research has reported sequential effects operating at the trial level. For example, a choice biases the interpretation of evidence in subsequent trials (Abrahamyan et al., 2016; Braun et al., 2018; Urai et al., 2019; Yu & Cohen, 2009). Similarly, a preliminary decision biases processing of additional post-choice evidence towards confirming the initial decision (Bronfman et al., 2015; Talluri et al., 2018; Rollwage et al., 2020), and decisions bias the strength-evaluation of pro-choice evidence that led to it (Jazayeri & Movshon, 2007; Luu & Stocker, 2018; Stocker & Simoncelli, 2008). Common to these studies is the assumption that these biases are triggered by the formation of a decision. Our findings, however, extend this notion by suggesting that a similar micro-level evidence integration bias operates during decision formation, prior to committing to a choice (see also Cheadle et al., 2014; Patai et al., 2020). In particular, we found that the best-fitting model was one in which the evidence is boosted for consistent evidence frames (this boost increasing with the number of consecutive consistent frames) and is reset to baseline when the first inconsistent piece of evidence is encountered. Despite its evidence distortion, we have shown that this mechanism has an adaptive function in the presence of integration noise. Since evidence samples that are consistent with their predecessors are more likely to carry stronger evidence in favor of the correct alternative, inflating their weight provides extra protection from the corrupting effect of accumulation noise, resulting in increased decision accuracy (Fig. 6). This ‘normativity hypothesis’ predicts that to the extent that consistency-based evidence integration is a controlled and adjustable strategy, consistency sensitivity effects will increase as a function of integration noise, for example, when one is performing a dual task or when working memory is loaded. We leave this interesting question for future studies (see Spitzer et al., 2017; for a parallel investigation in relating to selective-integration).

Our findings raise the intriguing hypothesis that confirmation biases are a form of consistency bias, whereby post-choice evidence inconsistent with pre-choice evidence is integrated less effectively than post-choice consistent evidence. Future studies should investigate whether, to what extent, and how, consistency and confirmation biases are related, by measuring both biases within participants and using a unified paradigm. Another interesting possibility is that the consistency of evidence supporting a decision might affect the extent of a confirmation bias. For example, choices that are based on more consistent evidence may likely be more prone to confirmation bias, for example, due to the mediating effect of decision confidence (Rollwage et al., 2020). Future studies could also beneficially examine whether and how consistency bias is related to a broad range of individual traits such as the need for cognitive closure (Kruglanski & Webster, 1996), need for consistency, political radicalism (Rollwage et al., 2018), and dogmatism, or to psychiatric conditions such as OCD (Cavedini et al., 2006). We believe our paradigm provides an important advantage over current confirmation bias paradigms by addressing these questions. In current confirmation bias paradigms, which probe one’s ability to revise initially-wrong decisions, validity might be jeopardized by demand characteristics (e.g., presenting oneself in a self-consistent manner). In contrast, the current approach eschews these concerns, since participants are not required to contradict or confirm their previous decisions.

In conclusion, our novel methods allowed us to validate critical aspects of the evidence accumulation process and to unravel the biases that affect it. Our findings contribute to a growing literature speaking to the notion that self-inflicted distortions of evidence are ironically adaptive in that they act to increase choice veracity by making it robust to noise. A critical next step is to study how these strategies are acquired and how they relate to puzzling behaviors such as confirmation bias, and to better characterize the environmental and psychological variables that affect strategy selection.

## Methods

### Experimental methods

#### Participants

Twenty-seven undergraduate from Tel-Aviv University (22 females) participated in Exp. 1^2^, thirty in Experiment 2 (22 females), and twenty-three in Exp. 3 (17 females), all of which reported having normal or corrected-to-normal vision. The participants received course credit in exchange for taking part in the experiments, as well as a bonus fee ranging from 15 – 25 ILS, which was determined by their task performance. The experiment was approved the ethics committee at TAU.

#### Stimuli

The stimuli were consisted of pairs of numerical values (Exp. 1 & 3) or bars (Exp. 2) which were presented simultaneously (see Fig. 1A-B for illustration), with a presentation rate of 2 Hz (Exp. 1 & 3) or 5 Hz (Exp. 2). Displays in Exp. 1 & 3 were generated by an Intel I7 personal computer attached to a 24’’ Asus 248qe monitor with a 144 Hz refresh rate, using 1920×1080 resolution graphics mode. Displays in Exp. 2 were generated by an Intel I3 personal computer attached to a 19’’ ViewSonic Graphics Series G90fB CRT monitor with a 60 Hz refresh rate, using 1024×768 resolution graphics mode. Responses were collected via the computer keyboard. Viewing distance was approximately 60 cm from the monitor.

#### Task and Design

Each trial in the experiments began with a fixation display which consisted of a black 0.2° × 0.2° fixation cross (+) that remained on the screen for 1s. Then, pairs of numerical values (Exp. 1 & 3) or bars (Exp. 2) were presented sequentially to the participants, who were asked to decide which of the sequences has a higher mean. The presentation in Exp. 1 & 2 was terminated by the participants’ response (free response protocol), while the presentation in Exp. 3 was terminated after eight samples (interrogation protocol). Responses were given by pressing the arrow keys (left/right arrow keys for the left/right sequences, respectively). In Exp. 2 & 3, after indicating the sequence with the higher mean, participants were also asked to indicate the confidence in their choice on a 1-6 scale.

#### Experimental Conditions

In all the experiments the samples were drawn from Gaussians distributions. Exp. 1 included two difficulty levels which were manipulated by varying the separation between the Gaussians: in the easy trials the means of the Gaussians were: μ_1_=52 vs. μ_2_=44, σ=10, and in the difficult trials the means were: μ_1_=52 vs. μ_2_=48, σ=10. Exp. 2 also consisted of two difficulty levels μ_1_ = 52.5 vs. μ_2_ = 47.5 and μ_1_ = 51.5 vs. μ_2_ = 48.5, as well as an orthogonal manipulation of the sequences variance: σ_1_ = 0.1167 vs. σ_2_ = 0.07. In Exp. 3 we orthogonally manipulated the difficulty and the consistency of the evidence. Difficulty was manipulated by increasing the separation between the Gaussians from μ_1_=52 vs. μ_2_=48, σ=10 (difficult trials) to μ_1_=52 vs. μ_2_=44, σ=10 (easy trials). Consistency was manipulated by sampling eight values from the high as well as from the low mean distributions. Then, to generate consistent trials we paired these values, so that in seven out of the eight pairs the stronger evidence was in favor of the higher mean distribution (see Fig. 4A, lower panel). To generate inconsistent trials, we shuffled the temporal order of the same values and repaired them, so that only in four out of the eight pairs the stronger evidence was in favor of the higher mean distribution (Fig. 4A, upper panel).

### Computational methods

#### Mixed effects models

The effect of accumulated evidence and stimulus-consistency on accuracy, RT confidence in Exp. 1 & 2 (Table 1) were estimated on a trial-by-trial basis using mixed model regression analyses. The regressions were implemented using Matlab’s ‘fitlme’ and ‘fitglme’ functions with participants serving as random effects and with a free covariance matrix. The fixed effects variables were: i) accumulated evidence - the sum of differences between the two streams of evidence at the moment of response, and ii) stimulus-consistency - the absolute value of the difference between the number of frames with evidence favoring the two alternatives, normalized by the length of trial. Both variables were normalized using z-score transformations.

Choices in Exp. 1 & 2 (coded as 1-correct and 0-error) were predicted using logistic regressions, which in Wilkinson notation was:

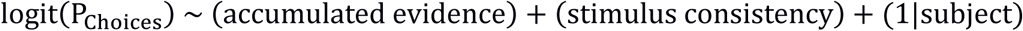

RTs (Exp. 1 & 2) and confidence (Exp. 2) were predicted using linear regressions, which in Wilkinson notation were:

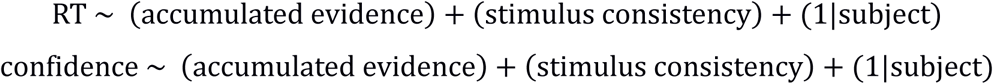

The exact same pattern of results reported in Table 1 was obtained if the accumulated evidence and stimulus-consistency were also included as random effects.

##### Model-free

The validity of the model-free method was tested by simulating 10,000 synthetic decisions using known (fixed or collapsing) boundaries, and examining the ability of the model-free method to accurately recover them. The values in all simulations (Fig. 2A-D) were sampled from *X~N(52, 15^2^)* and *Y~N(46, 15^2^)*. The decision process in Fig. 2A-B was based on Eq. 1 with either fixed (Fig. 2A) or collapsing boundary (Fig. 2B). The fixed-boundary was characterized by a single boundary parameter (c=50), and the collapsing-boundary was characterized by four parameters: intercept, shape, scale and asymptote parameters (a=100, k=3, λ=4, a’=20; see Computational Methods and Hawkins et al., 2015). The decision process in Fig. 2C-D was based on the value and difference cut-off heuristics with cut-offs of 70 and 20, respectively (see Computational Methods and Glickman & Usher, 2019).

The decision boundaries in Fig. 2A-D were extracted by applying the LDA algorithm (Fisher, 1936; McLachlan, 2004) to the integrated evidence excluding internal noise (i.e., Y(t), see Eq. 2) of each frame of all 10,000 synthetic trials. For each frame, each trial was classified as one of the following categories: choose alternative-A, choose alternative-B or continue sampling. Then, using the LDA, we extracted the planes that optimize the separation between different classes for each frame. We assumed that the upper and lower boundaries are symmetrical and therefore averaged both.

As mentioned in the main text, the internal noise causes an unavoidable overlap between the different classes. This overlap impairs the LDA ability to correctly extract the decision boundary, and is particularly evident in slow trials due to the accumulation of internal noise across time (Teodorescu, Moran, & Usher, 2016). Thus, in order to increase the robustness of the model-free method to internal noise, we constrained the boundary extraction of each frame by previous ones. To this end, we extracted the boundary based on two features: *t* and *Y*(*t*), 1 ≤ *t* ≤ *n*, as illustrated in the table below:

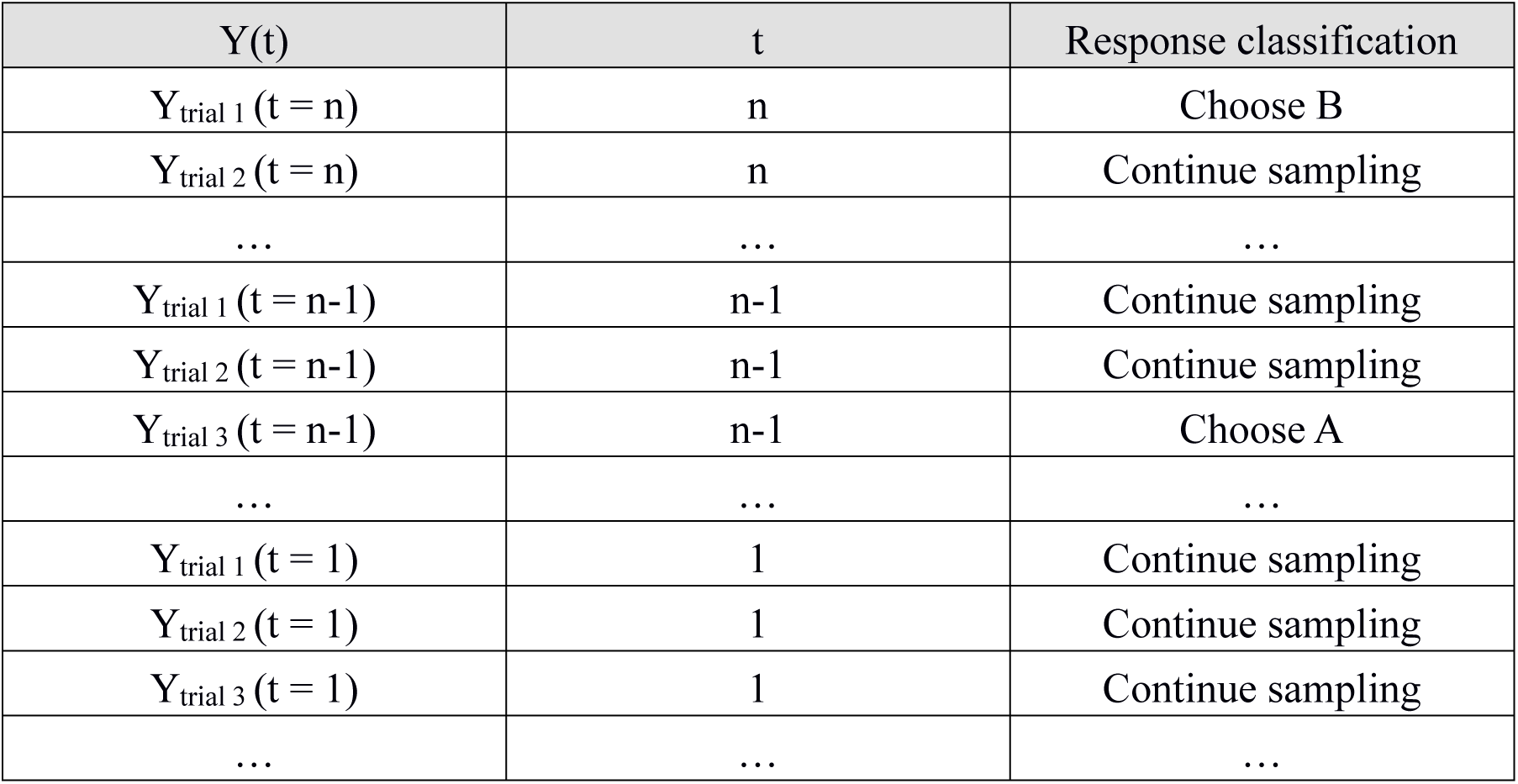

The LDA algorithm provide linear functions that separate the different classes from each other. To obtain the value of the boundary for the n-th frame, we computed the value of the separating linear functions for this frame.

### Modelling Details

#### Integration to boundary models

We examined several integration-to-boundary models, all of them assumed integration of evidence based on the following formula:

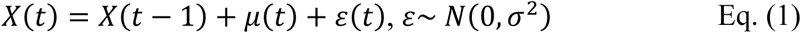

where *X(t)* is the accumulated differences at time *t*, *ε*(*t*) is a random Gaussian noise, which is independent from the evidence-sampling noise. The term *μ*(*t*) varied between the different models as follows:

i. **Full integration**

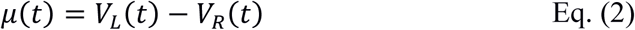

where *V*_*L*_(*t*) and *V*_*R*_(*t*) are the samples drawn from the left and right distributions at time t, respectively (note that *V*_*L*_(*t*) and *V*_*R*_(*t*) include the sampling noise).
ii. **Stimulus consistency**

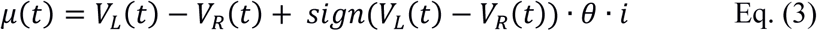

where *θ* is a free parameter representing the enhancement given to pieces of evidence that are consistent with previous ones, and *i* counts the run of consistent values (starting at 0). For example, if the differences between the values are: 15, 20, 8, −15 and −25, then *i*_*t*=1_ = 0, *i*_*t*=2_ = 1, *i*_*t*=3_ = 2, *i*_*t*=4_ = 0 and *i*_*t*=5_ = 1.
iii. **Preference consistency**

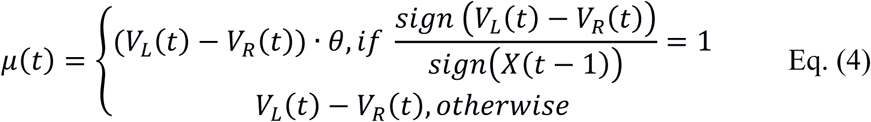

where *θ* is a free parameter representing the enhancement given to pieces of evidence consistent with the total accumulated evidence at time *t* − 1.
iv. **Selective integration**

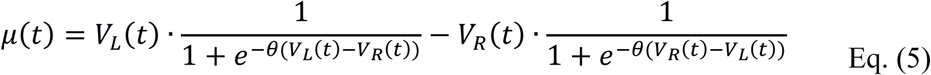

where *θ* is a free parameter affecting the magnitude of the selective gating (see Glickman et al., 2018).

All the integration models in Exp. 1 & 2 assume integration to a collapsing-boundary (see Glickman & Usher, 2019), modeled using a Weibull cumulative distribution function (Hawkins et al., 2015):

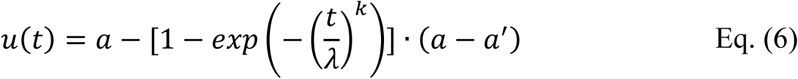

where *±u(t)* are the upper/lower thresholds at time *t*, *a*/*a′* are the initial (intercept) and asymptotic values of the boundary, respectively, and *λ/k* are the scale and shape parameters of the Weibull function, respectively.

In Exp. 3 we used an interrogation paradigm, in which the probability of choosing each alternative was calculated using an exponential version of Luce’s choice rule (Luce, 1959):

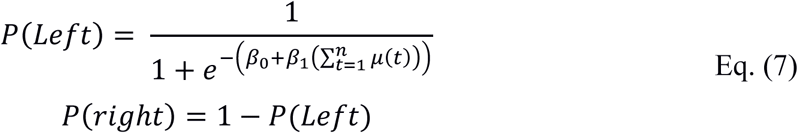

where *β*_1_ indicates the sensitivity of the model to the accumulated evidence, with an intercept of *β*_0_.

In addition, we examined whether the participants in Exp. 3 showed a recency bias, as reported in several previous studies which used an interrogation paradigm (Tsetsos et al., 2012; 2016). To this end, for each participant, we performed a temporal logistic regression analysis, in which we predicted the response of each trial based on the differences between *V*_*L*_ and *V*_*R*_ at each frame, ranked by their temporal order. As shown in Figure 4B, the mean weight of samples 5-8 was higher than that of samples 1-4 (*t*(21) = 9.97, *p* <. 001), indicating a recency bias. This motivated us to include a leak term that controls the extent to which earlier values are given less weight (Usher & McClelland, 2001). Thus, Eq. 7 was extended to the following form:

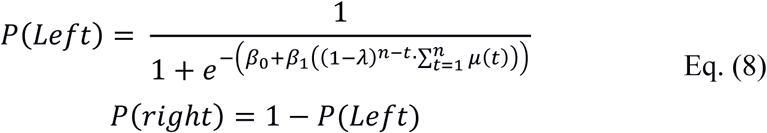

where *λ* is the leak term.

#### Non integration to boundary models

We examine three models that did not assume integration of evidence decision boundary (Fig. 1C-D and Fig. 2C-D). The first model is the value-cutoff heuristic, which assumes that observers choose based on the detection of a single high-value sample. For example, if a participant uses a cut-off value of 70, then s/he will choose the sequences in which a value higher than 70 first appears. The second heuristic is the difference-cutoff heuristic, which assumes that observers choose based on the first frame in which the difference between the numbers exceeds a predetermined threshold (Glickman & Usher, 2019). In addition to these two heuristics, we examined a third model which we labeled a random-timer model. This model assumes integration of evidence based on Eq. 1, however, response-time is determined by an exogenous process. The value cut-off heuristic was simulated using thresholds of 70 (Exp. 1, Fig. 1C), 80 (Exp. 2, Fig. 1D) and 70 (model-free, Fig. 2C). The difference cut-off heuristic was simulated using thresholds of 20 (Exp. 1, Fig. 1C), 25 (Exp. 2, Fig. 1D) and 20 (model-free, Fig. 2D). Response-times of the random timer model were sampled from an ex-Gaussian distribution with μ = 3, σ = 0.5 and λ = 2/3 (Exp. 1, Fig. 1C) and μ = 6, σ = 1 and λ = 3 (Exp. 2, Fig. 2C). These values were chosen because they provided accuracy and response-times similar to the ones observed in the data of Exp. 1 & 2.

#### Optimization procedure

The free parameters of the computational models were fitted to the data (choices and decision-times) of each participant in Exp. 1 & 2 separately, using maximum likelihood estimation (MLE). For each trial, we simulated the different models 1,000 times for a given set of proposal parameters and calculated the proportion of trials in which the model choice and decision-time matched the empirical data. Denoting the proportion of match between the simulated and empirical data by *p*_*i*_, we maximized the likelihood function *L*(*D*|*θ*) of the data (*D*) given a set of proposal parameters (*θ*), by:

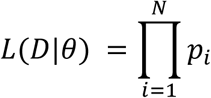

To find the best set of proposal parameters we first used an adaptive grid search algorithm (see Tavares, Perona, & Rangel, 2017 for details) and then used the three best sets of proposal parameters as starting points to a Simplex minimization routine (Nelder & Mead, 1965).

## Acknowledgments

This research was supported by grants to Marius Usher from the United States-Israel Binational Science Foundation (CNCRS = 2014612). We wish to thank Dr. Douglas Lee for his critical reading of the manuscript and helpful comments.

## Supplementary Information

### A) Behavioral signature of integration to boundary

We compared integration and a non-integration models by using the method described in Glickman & Usher (2019). Each model was simulated 100,000 times and the mean accumulated evidence was plotted as a function of RT (see black, green and magenta dashed lines in Fig. 1C-D). Next, for each participant in Exp. 1 & 2, the accumulated evidence excluding internal noise (both of correct and incorrect responses) was plotted as a function of response-time (see Fig. S1A-B). A mixed-effect linear regression models were fitted to the data of Exp. 1 & 2, with RT and participants serving as random effects and with a free covariance matrix (see thin gray and blue lines in Fig. 1C-D). The model in Wilkinson notation was:

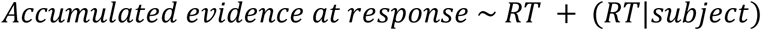

**Fig. S1.**
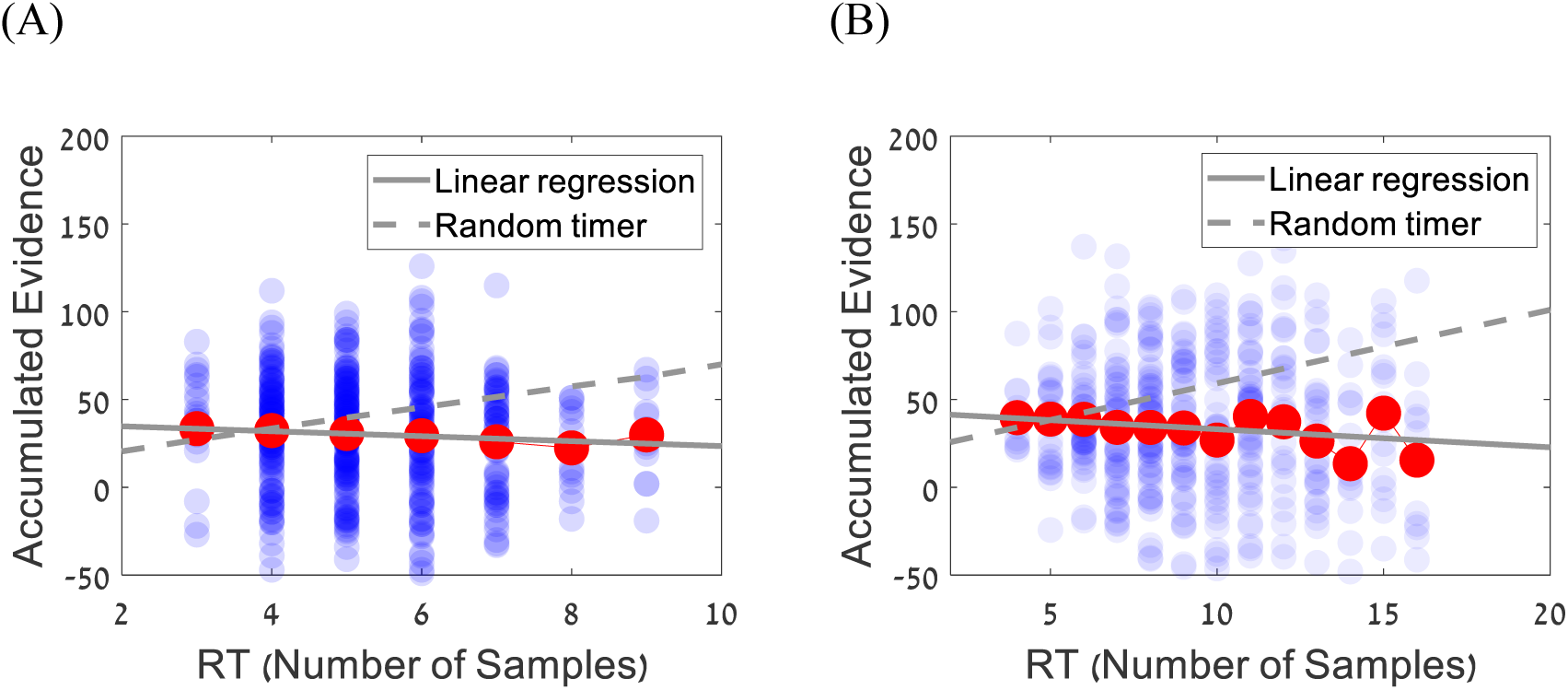
Scatter-plot showing the accumulated evidence in individual trials (blue circles) as a function of decision time of a representative participant in Exp. 1 (A) and Exp. 2 (B). Red circles correspond to the average accumulated evidence at each RT, the grey solid line corresponds to linear regression fitted to the data, and the dashed gray line corresponds to the prediction of the random timer model (see Computational Methods).

### B) Regression analysis - Largest temporal cluster of the evidence

**Table S1.** Beta coefficients for predicting Accuracy, RT and confidence in Exp. 1&2.

|  | **Exp. 1**<br>**(Numerical Stimuli)** |  | **Exp. 2**<br>**(Perceptual Stimuli)** |  |  |
| --- | --- | --- | --- | --- | --- |
|  | **Accuracy** | **RT** | **Accuracy** | **RT** | **Confidence** |
| **Evidence** | $\beta = 1.79^{***}$<br>$t = 32.04$ | $\beta = -0.29^{***}$<br>$t = -35.57$ | $\beta = 0.53^{***}$<br>$t = 17.58$ | $\beta = -0.19^{***}$<br>$t = -28.12$ | $\beta = 0.18^{***}$<br>$t = 20.95$ |
| **Stimulus Consistency (LTC)** | $\beta = 0.20^{***}$<br>$t = 4.68$ | $\beta = -0.14^{***}$<br>$t = -16.60$ | $\beta = 0.97^{***}$<br>$t = 27.43$ | $\beta = -0.16^{***}$<br>$t = -24.25$ | $\beta = 0.10^{***}$<br>$t = 10.99$ |

\*\*\* $p < .001$

The choice-accuracy, RT and confidence in Exp. 1 & 2 were analyzed using mixed model regression analyses (logistic for accuracy and linear for RT and confidence), using the accumulated evidence and normalized stimulus-consistency as fixed factors and participants as random intercepts. The analysis was similar to one reported in Table 1 (see also Computational models), except here we used the more complex measure of stimulus-consistency – larger temporal cluster of the evidence (LTC), in which for each trial we computed the largest temporal cluster that goes with the evidence and divide it by the number of samples (e.g., if the evidence are: 4, −5, 3, 4, 5, 2, −2, −6, then the LTC measure will be equal to: 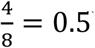). The results (Table S1), are fully consistent with the ones obtained using the simpler consistency measure (Difference in Evidence-Directions), and show that LTC improves accuracy and confidence and reduces RT, even while controlling for the accumulated evidence.

### C) Model based vs. Model free - Exp. 1

**Fig. S2.**
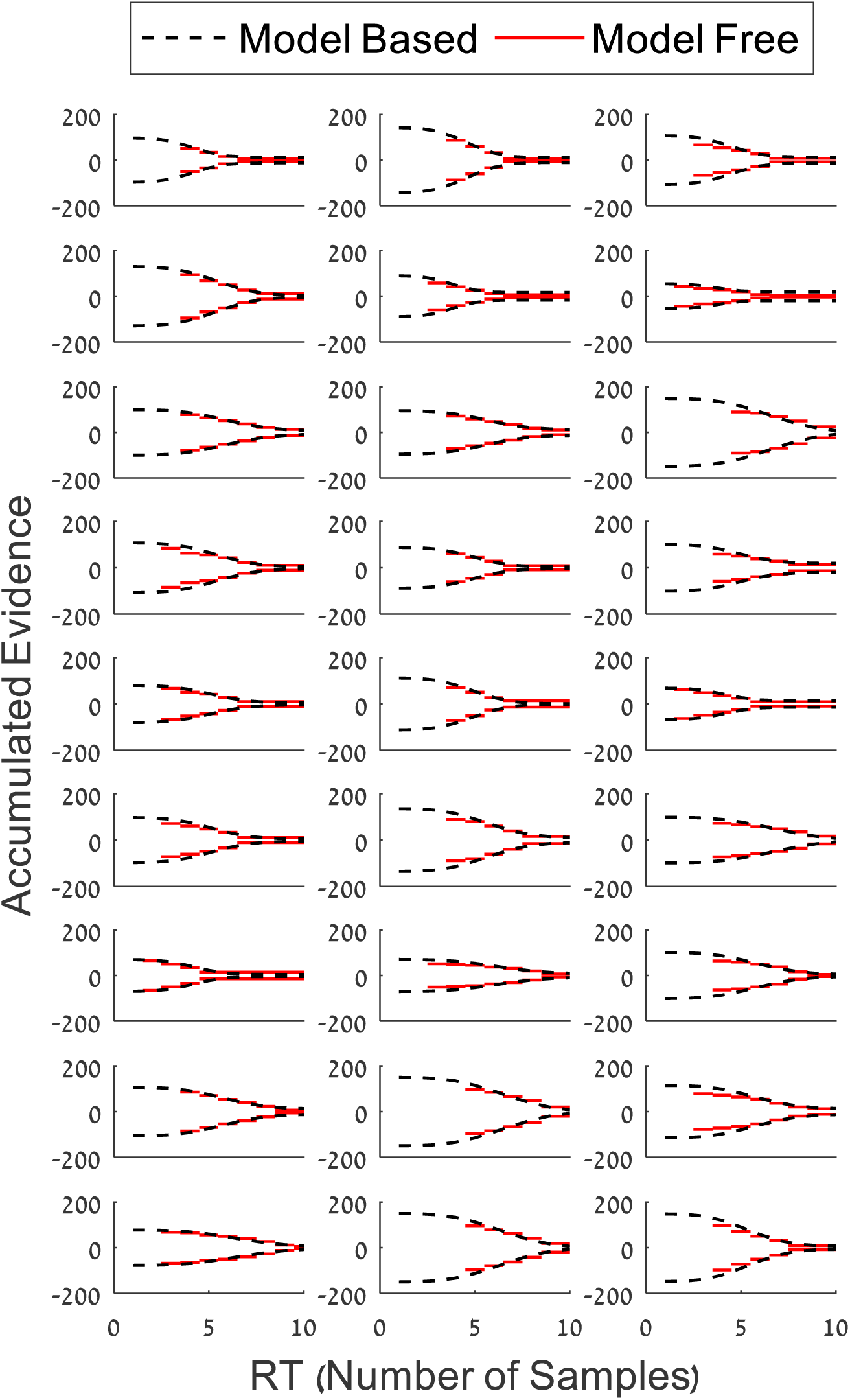
Decision Classification Boundaries (BCD) of the participants in Exp. 1. Black dashed lines correspond to the model-based DCB, and the red ones correspond to the model-free DCB.

#### Model based vs. Model free - Exp. 2

**Fig. S3.**
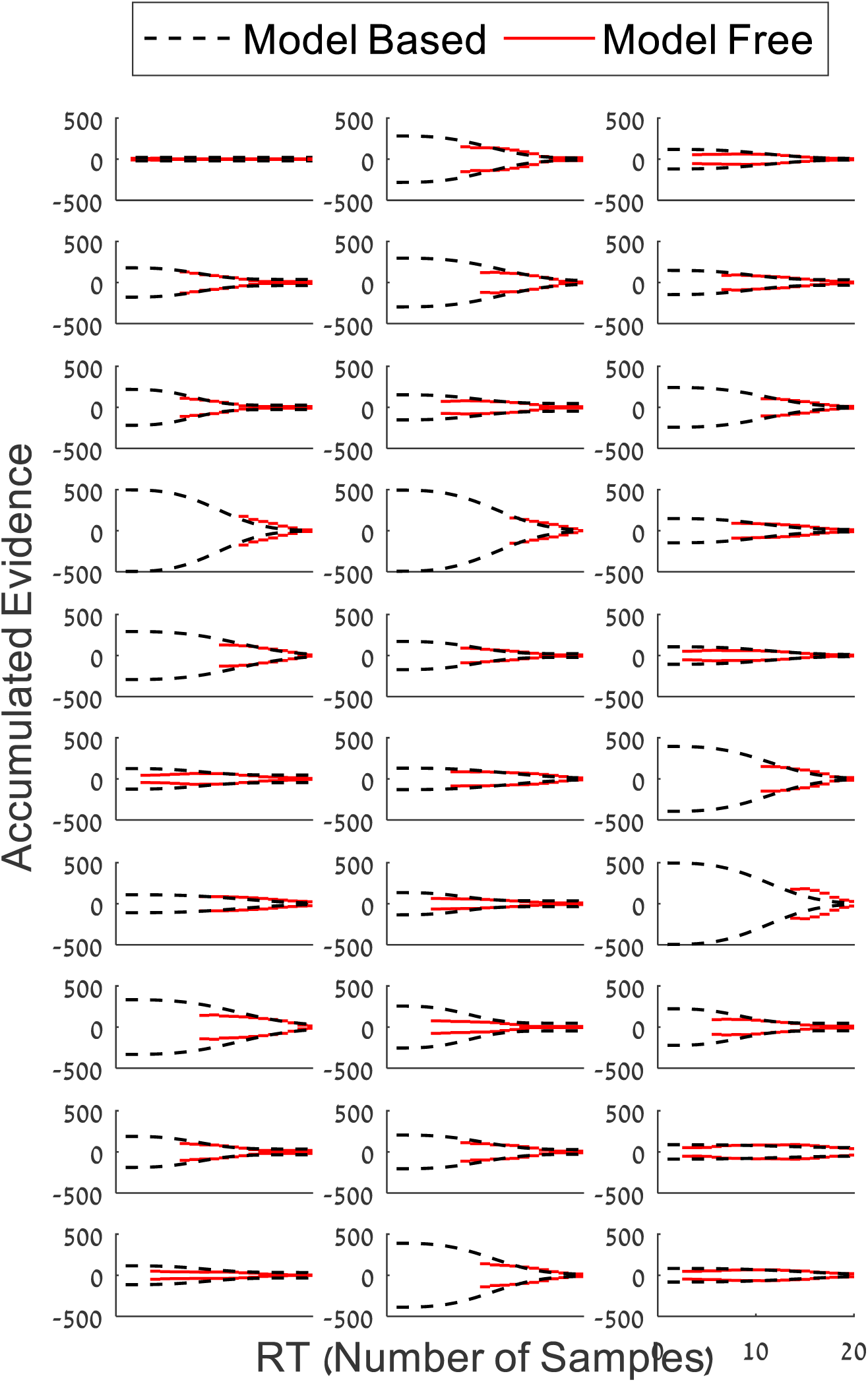
Decision Classification Boundaries (BCD) of the participants in Exp. 2. Black dashed lines correspond to the model-based DCB, and the red ones correspond to the model-free DCB.

### D) DCB modulation by stimulus consistency – Exp. 1

**Fig. S4.**
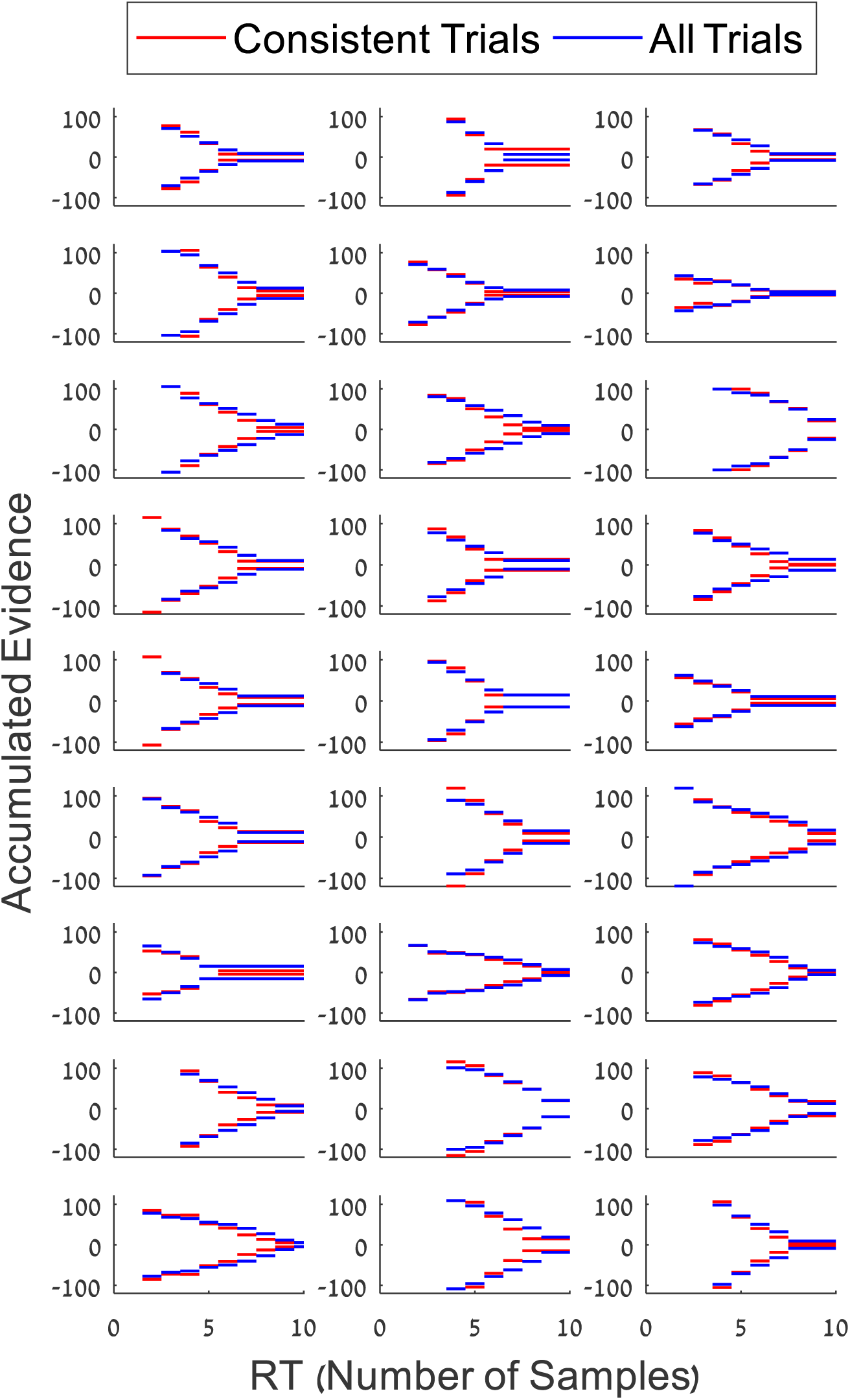
Decision boundaries for all trials (blue) and high consistency trials (red) of all participants in Exp. 1 (numerical stimuli). The boundaries were extracted using the model-free method.

#### DCB modulation by stimulus consistency – Exp. 2

**Fig. S5.**
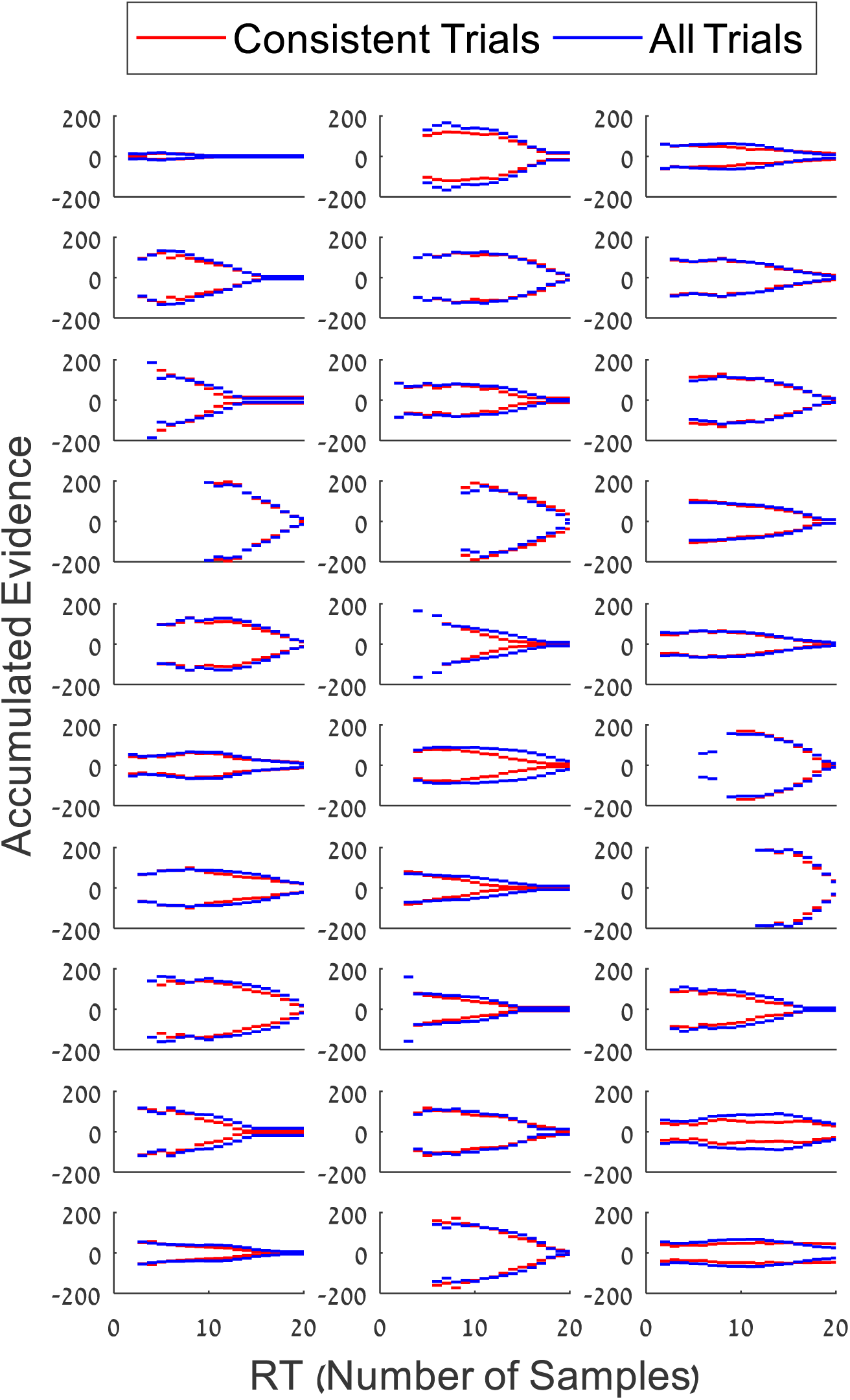
Decision boundaries for all trials (blue) and high consistency trials (red) of all participants in Exp. 2 (numerical stimuli). The boundaries were extracted using the model-free method.

#### DCB modulation by stimulus consistency – Quantitative comparison

In order to quantitatively compare the Decision-Classification Boundary (DCB) of high consistency trials with that of all trials, we computed the difference of the area below the boundary (i.e., the area between the upper boundary and the x-axis) for the two conditions and normalized it by dividing by the sum of the areas:

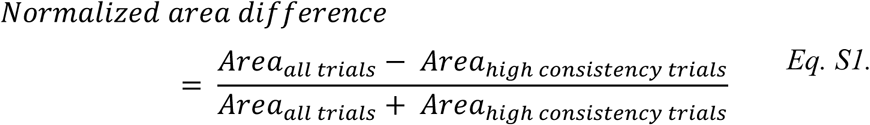

If participants require less evidence to reach a decision in high consistency trials, then the decision boundary of these trials should be lower than the decision boundary of all trials, resulting in normalized area difference higher than 0. A single-sample *t*-tests (against 0) confirmed that this was the case in Exp. 1, *t*(26) = 2.67, *p* =. 01 (Fig. S6A), as well as in Exp. 2, *t*(26) = 3.91, *p* <. 001 (Fig. S6B).

**Fig. S6.**
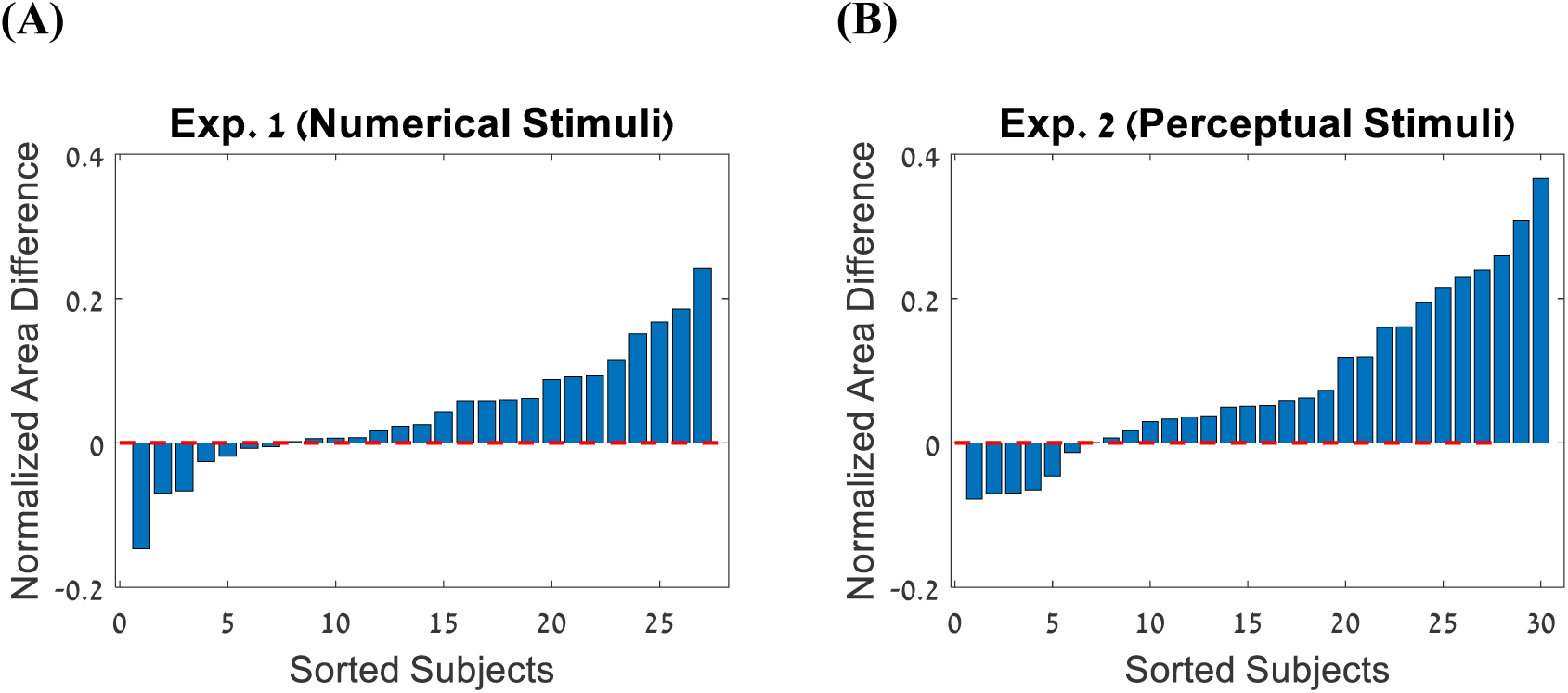
(A) Normalized area differences of the participants in (A) Exp. 1 and (B) Exp. 2.

We then tested that this DCB separation (consistent trials vs. all trials) takes place in synthetic data for which choices are generated by the stimulus-consistency model, but not when generating data from the full-integration model. Toward this aim, we simulated the full-integration and the stimulus-consistency models using sequences with the same statistical properties as the ones in Exp. 1 & 2, and with the best fitted parameters of each participant. The DCB of each synthetic participant was extracted using the same procedure as the one used for the empirical data. Fig. S7A-B shows that the DCB separation effect is only predicted by the stimulus-consistency model, *t*(26) = 2.45, *p* =. 02 (Exp. 1) and *t*(26) = 2.77, *p* =. 009 (Exp. 2), and not by the full-integration model, *t*(26) = 0.75 *p* =. 45 (Exp. 1) and *t*(26) = −0.61, *p* =. 54 (Exp. 2).

**Fig. S7.**
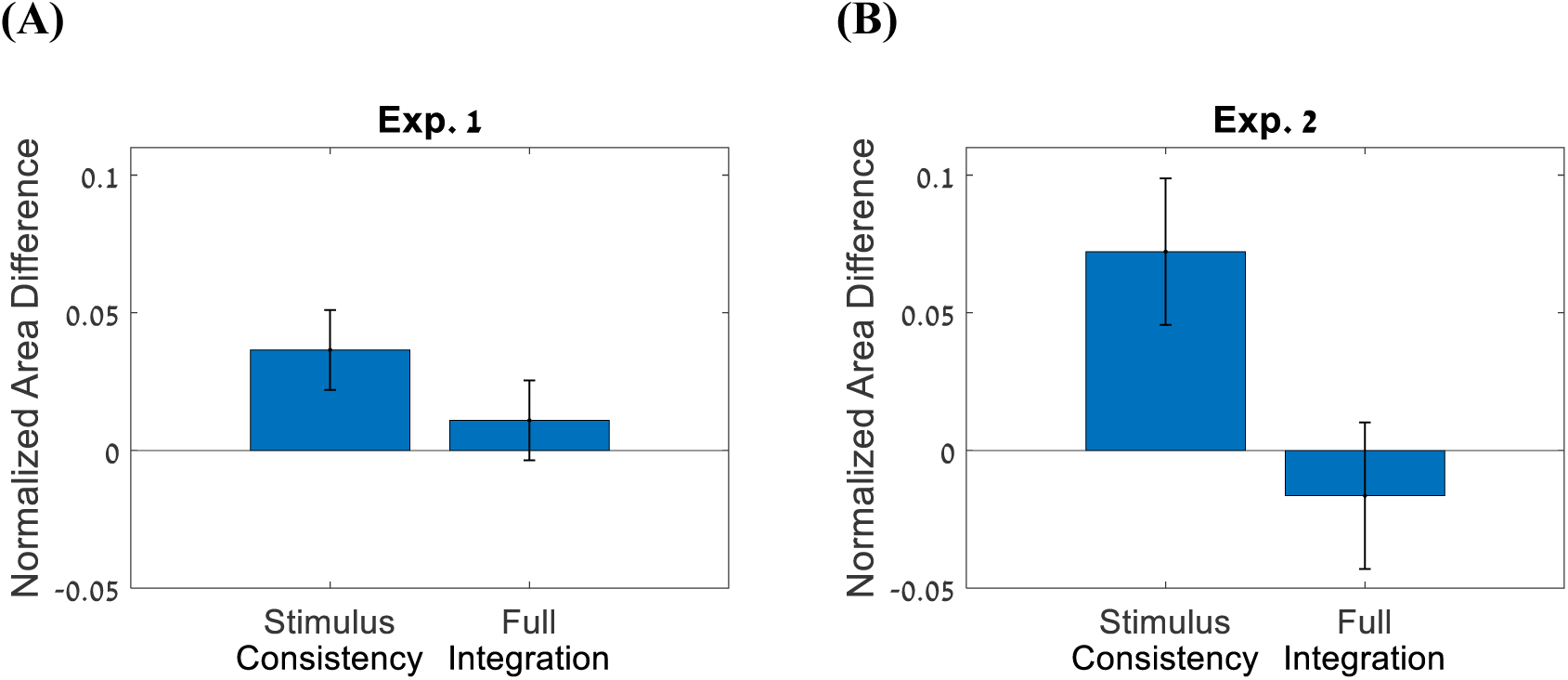
Mean normalized area difference of the stimulus consistency and the full integration models in (A) Exp. 1 and (B) Exp. 2. Error bars correspond to standard-error of the mean.

### E) Temporal clustering effects

In Exp. 3 we generated consistent and inconsistent trials (Fig. 4A), by manipulating the difference between the number of frames with evidence favoring the two alternatives. This manipulation, however, is correlated with the number of frequent winners, which modulates choices in the selective-integration (SI) model (Tsetsos et al., 2016). Thus, although the stimulus-consistency (SC) model provided better fit for the data (Fig. 5), the consistency effect in Exp-3 can also be accounted for by the SI model (Fig. 4C). To distinguish between the two models, we used another measure of consistency – LTC (larger temporal cluster of the evidence, see Table S1). For each participant, we calculated the LTC measure of each trial, and divided the trials into high/low LTC trials by performing a median split. Then, to quantify the sensitivity of the participants to temporal clustering, we computed the difference in mean accuracy of high and low LTC trials (accuracy_high LTC_ – accuracy_low LTC_). This difference should be higher for a participant whose bias mechanism is modulated by the consistency of each frame with previous ones, as presenting the positive evidence in an as large cluster as possible enhances the overall integrated evidence.

Fig. S8A shows the differences in goodness of fit (*deviance =-2∙LogLikelihood*) between the SI and the SC models (positive values indicate a better fit of the SC model). As can be seen, the SC model provides better fit than the SI-model (for all except four participants). Fig. S8B shows the correlation between differences in the goodness of fit of the models and the effect of temporal clustering (each point represents a single participant). As can be seen, the SC model provide a better fit for the data, specifically for participants who showed higher sensitivity to temporal clustering.

**Fig. S8.**
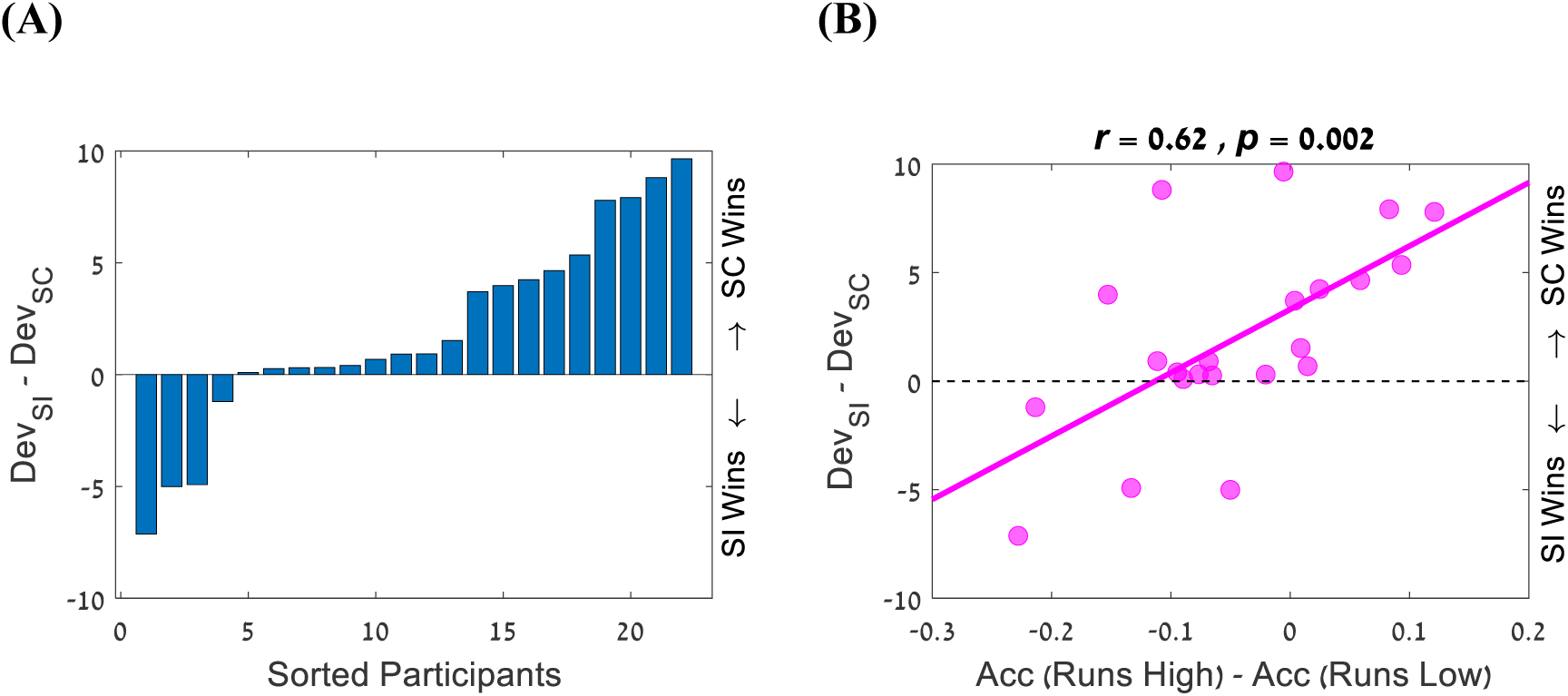
(A) Comparison between the SC and SI models across the 22 participants in Exp. 3. (B) Correlation between differences in goodness of fit of the SI and the SC models and the effect of temporal clustering; each point represents a single participant.

### F) Confidence-resolution

In order to account for the differences in confidence resolution in Exp. 3 (Fig 4D), we simulated the stimulus-consistency model using pairs of consistent and inconsistent trials, that were created using the same generating distributions as in Exp. 3 (i.e., with the same evidence content for each pair). Fig. S9A shows 25 pairs of trials (blue – consistent vs. red – inconsistent), which were simulated using the mean fitted parameters: *θ*_*consistency parameter*_ = 1.75 and *λ* = 0.21. The stimulus-consistency model predicts that the bias in favor of consistent evidence would increase the accumulated evidence of consistent compared with inconsistent trials (even though both types of trials have the same evidence). Fig. S9B shows the distributions of accumulated evidence of 100,000 consistent and inconsistent trials simulated using the stimulus-consistency model (using the mean internal noise obtained in the empirical data, *β* =. 05). The mean distance from the criterion of correct consistent trials is higher than for correct inconsistent trials *t* = 3.55, *p* <. 001. For inconsistent trials, however, the differences in the mean distances from the criterion of the incorrect consistent and incorrect inconsistent trials did not reach statistical, *t* = 0.25, *p* =. 79. Based on the distributions presented in Fig. S9B we predicted the mean confidence response for each condition of the stimulus consistency. To this end, we computed the absolute mean value of the accumulated evidence (i.e., distance from 0) of the correct and incorrect responses separately for consistent and inconsistent response. Then, using linear regression we mapped these values to the mean confidence level of each condition (Fig. S9C, dashed lines).

**Fig. S9.**
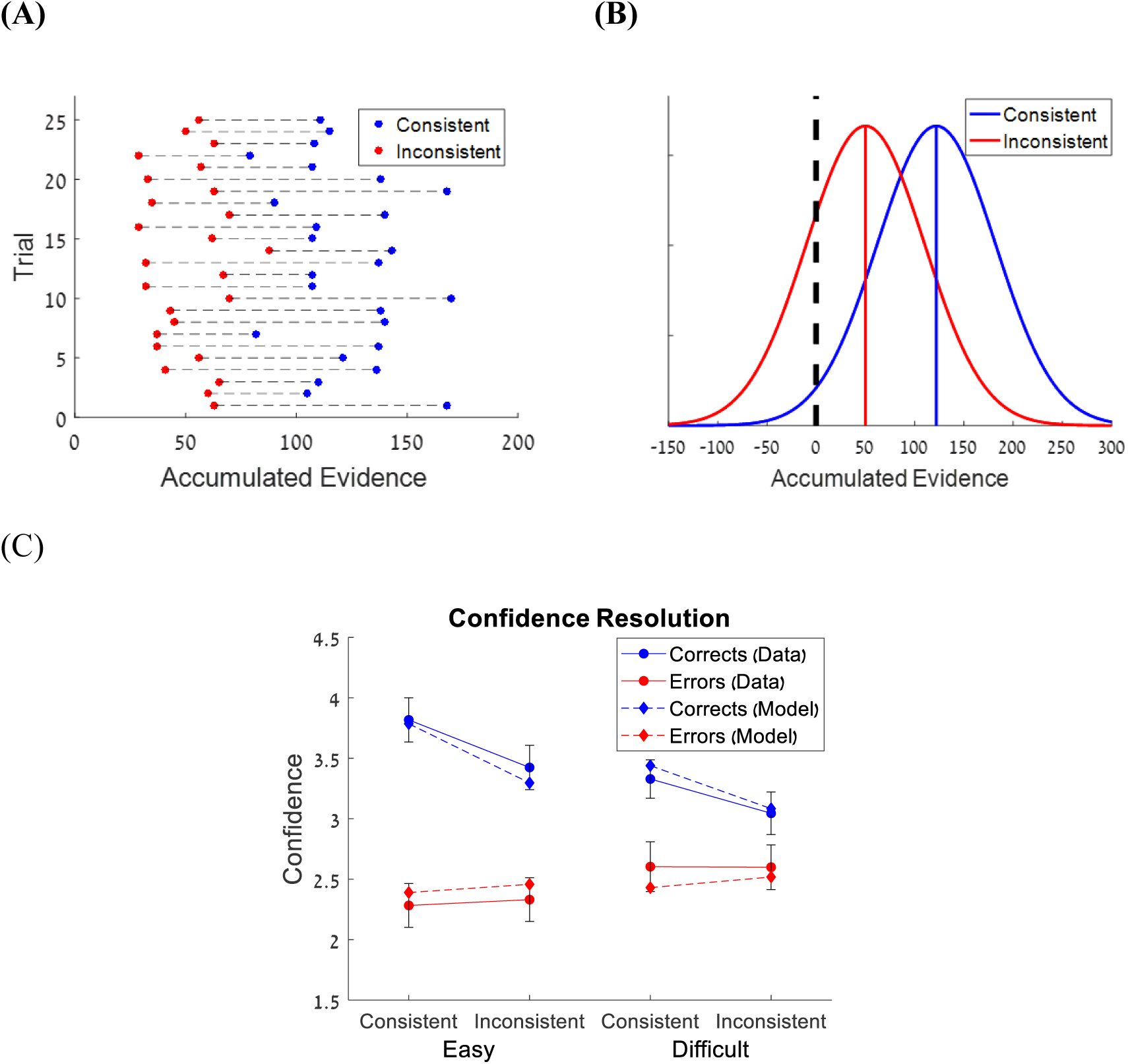
Confidence-resolution in Exp. 3. (A) Twenty-five pairs of trials (consistent and inconsistent) were simulated using the same generating distributions as in Exp. 3 (without internal noise; blue – consistent trials, red - inconsistent). The stimulus-consistency model predicts that the bias in favor of consistent evidence would increase the accumulated evidence of consistent trials compared with inconsistent ones. (B) The distributions of accumulated evidence of consistent (blue) and inconsistent (red) trials simulated using the stimulus-consistency model (with internal noise). Solid blue and red vertical bars correspond to the mean of each distribution. (C) Confidence as a function of difficulty and consistency for correct (blue lines) and incorrect (red lines) responses. Data are shown with solid lines and circle symbols and model predictions are shown with dashed lines and diamond symbols. Error bars are within-subjects SE.

## Footnotes

1 Since we are looking at normative considerations, those simulations exclude integration-leak; adding it to all models does not change any of the results.

2 The data of Exp. 1 was taken from our previously published paper (Glickman & Usher, 2019).

